# Induction of immortal-like and functional CAR-T cells by defined factors

**DOI:** 10.1101/2023.11.06.565785

**Authors:** Lixia Wang, Gang Jin, Qiuping Zhou, Yanyan Liu, Xiaocui Zhao, Zhuoyang Li, Na Yin, Min Peng

**Affiliations:** Department of Basic Medical Sciences, School of Medicine, Tsinghua University, Beijing 100084, China; Institute for Immunology, Tsinghua University, Beijing 100084, China; Tsinghua-Peking Center for Life Sciences, Beijing 100084, China; Beijing Key Laboratory for Immunological Research on Chronic Diseases, Tsinghua University, Beijing 100084, China

## Abstract

Long-term antitumor efficacy of chimeric antigen receptor (CAR)-T cells depends on their functional persistence in vivo. T cells with stem-like property show better persistence, but factors conferring T cells bona fide stemness remain to be determined. Here, we demonstrate the induction of CAR-T cells into an immortal-like and functional state, termed TIF. The induction of CAR-TIF cells depends on repression of two factors: BCOR and ZC3H12A, and requires antigen or CAR tonic signaling. The reprogrammed CAR-TIF cells possess almost infinite stemness resembling induced pluripotent stem cells but retain functionality of mature T cells, which exhibit superior antitumor effect. CAR-TIF cells enter a metabolically dormant state after elimination of target cells, which persist in vivo with a saturable niche and mediate memory protection. CAR-TIF cells represent a novel state of T cells with unprecedented stemness conferring long-term functional persistence of CAR-T cells in vivo, which have broad potentials in T cell therapies.

## Main

Adoptive T cell therapy (ACT) is a pillar of cancer immunotherapy ^1, 2, 3^. Long-term persistence of adoptively transferred T cells is critical for sustained therapeutic efficacy ^4^,5. In human patients and mouse models, T cells with memory, stem-like or precursor phenotype confer better therapeutic outcomes ^6, 7, 8, 9, 10^, which makes induction of stem-like properties in endogenous or adoptively transferred T cells a major goal of cancer immunotherapy ^4, 5^. Due to their paramount importance in immunity and immunotherapy, stem-like T cells such as central memory T (TCM) cells ^11^, memory stem T (TSCM) cells ^6, 7^ and precursor exhausted T (TPEX) cells ^12, 13, 14, 15, 16^ have been extensively studied. However, it remains unknown how to efficiently produce large numbers of T cells with bona fide stemness for ACT.

The obstacles that make production of large amount of stem-like T cells difficult, if not possible, root in the fundamental nature T cell response and intrinsic property of stem cells. A canonical feature of T cell response is massive contraction after clonal expansion ^17^, which leaves a few cells mediating memory protection ^18^. Thus, maintaining large quantities of antigen-specific T cells after peak response is against the natural kinetics of T cell response. Similarly, cells with self-renewing capacity such as stem and precursor cells are essentially rare in adult animals ^19^, thus generation and maintaining large quantities of stem-like cells does not fit the paucity nature of stem cells. Furthermore, stemness is usually mutually exclusive with functionalities ^20^. Conferring stemness to mature cells via cellular reprogramming, such as induction of induced pluripotent stem cells (iPSCs) ^21^, is at the cost of losing identity and function of cell-of-origin, which is not suitable for cellular reprogramming intended for disease therapy ^20^. How to reprogram differentiated mature cells to gain bona fide stemness but at the same time retain their identity and physiological function remains elusive ^20^.

## Results

### Induction of immortal-like and functional CD19 CAR-T (CAR19TIF) cells by repressing ZC3H12A and BCOR

After CD19 CAR-T (CAR19T) cells therapy of B cell malignancies, the majority of patients eventually relapsed due to antigen loss or poor persistence of CAR-T cells ^22^. Antigen loss can be circumvented by targeting other B cell antigens (e.g. CD22 and BCMA), which makes poor persistence of CAR-T cells a major obstacle for durable efficacy. CAR19T cells recognize both normal and malignant B cells. Although mature B cells are eliminated during primary response, hematopoietic stem cells (HSCs) continuously produce CD19^+^ progenitor cells that cannot be completely cleared. Indeed, rebound of CD19^+^ normal B cells in peripheral is an early sign of relapse after treatment with CAR19T cells ^23^. To mimic the clinical situation where infused CAR19T cells recognize endogenous B cells, we used a published CD19 CAR targeting mouse CD19 (mCD19) ^24^ (Fig. 1a). T cells transduced with this mCD19 CAR could eliminate both endogenous and malignant B cells when combined with lymphodepleting conditioning ^24^. However, without conditioning, CAR19T cells could not expand nor kill target cells in immunocompetent mice ^24^, which explains why chemotherapeutic conditioning is an essential part of CAR-T cell therapy in clinic. We confirmed that CAR19T cells did not expand nor kill CD19^+^ cells in immunocompetent mice without conditioning (Extended Data Fig. 1a-c). To develop a conditioning-free and durable CAR-T cell therapy in immunocompetent hosts, all ACT in this study were performed without any conditioning regimen such as chemotherapeutic treatment (e.g. cyclophosphamide and/or fludarabine), total body irradiation, vaccination, cytokine infusion, et al.

**Fig. 1.**
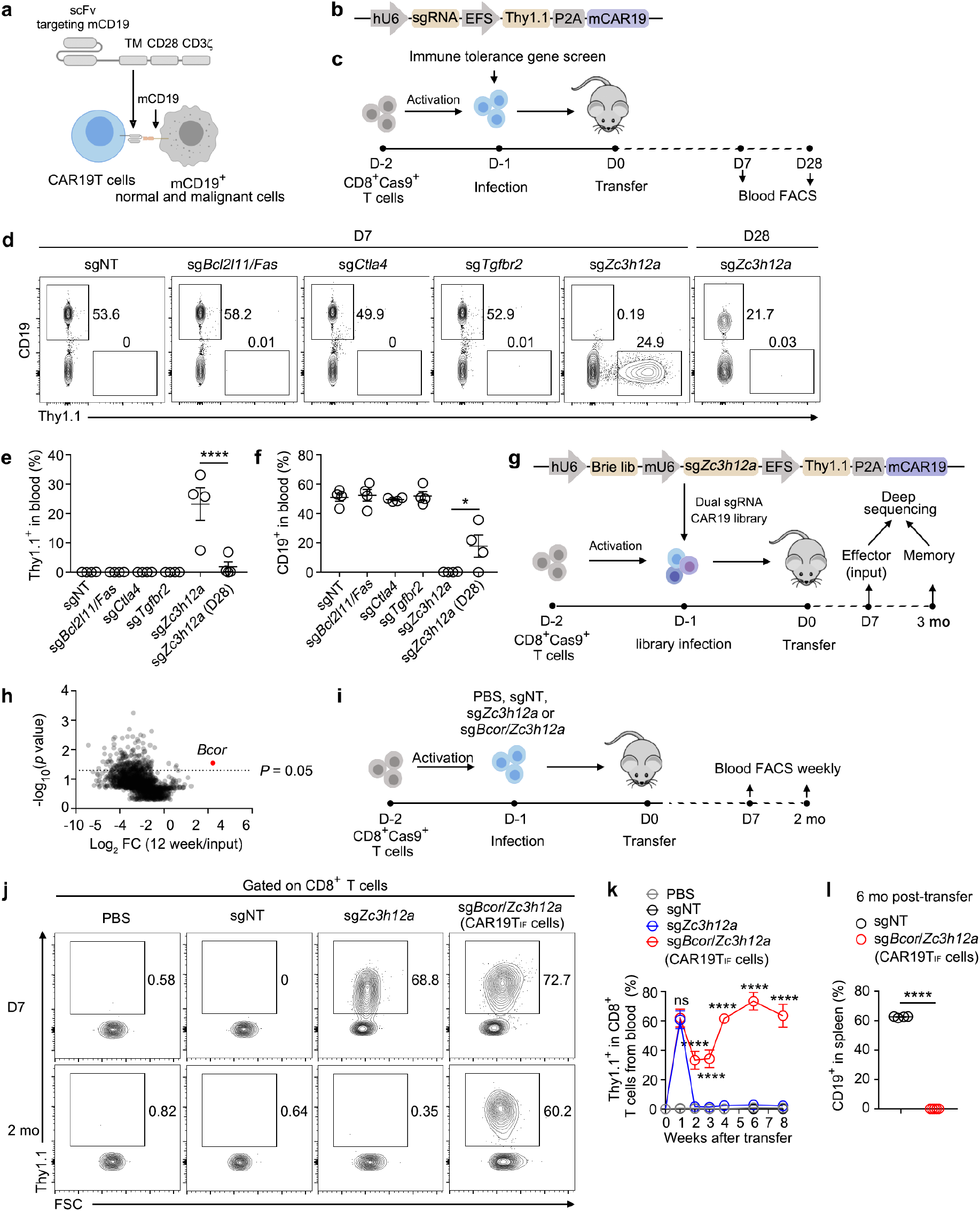
Ablation of ZC3H12A and BCOR generates immortal-like and functional CAR19TIF cells. **a,** The composition of anti-mCD19 CAR. **b**, Vector design of pMSCV- sgRNA-CAR19. Thy1.1 is co-expressed with CAR at protein level via P2A, serving as a marker of CAR expression. **c**, Experimental design. CD8^+^Cas9^+^ T cells were activated by anti-CD3/anti-CD28 (1 μg/ml), infected with retrovirus expressing CAR19 with sgRNA targeting indicated genes. One million of CD8^+^Thy1.1^+^ CAR19T cells were transferred into B6 mice. CAR19T cells and B cells from peripheral blood were monitored by flow cytometry. **d-f**, Representative plots (**d**) and statistical analysis (**e,f**) of Thy1.1^+^ CAR19T cells and CD19^+^ B cells among single live cells from peripheral blood 7 days and 28 days post-transfer are shown (n = 4 mice for each group). **g**, Experimental design for genome-wide dual sgRNA screening for gene whose loss of function confers persistence to ZC3H12A-deficient CAR19T cells. Indicated sgRNA library was used to infect activated CD8^+^ T cells, and 150 million CAR19T cells were transferred into 50 B6 mice (3 million per mouse). Seven days and 3 months later, Thy1.1^+^ CAR19T cells were sorted from spleen for sgRNA analysis. **h**, Screening result. **i**, Experimental design for validation of the screening hit BCOR. **j,k**, Representative plots (**j**) and statistical analysis (**k**) of Thy1.1^+^ CAR19T cells and CD19^+^ B cells among single live cells from peripheral blood of mice transferred with CAR19TIF cells expressing indicated sgRNAs are shown (n = 3 mice in PBS group, n = 6 mice in sgNT group, n = 5 mice in sg*Zc3h12a* group, n = 5 mice in sg*Bcor*/*Zc3h12a* group). **l**, Percentages of CD19^+^ B cells from spleen of mice 6 months after transferring CAR19TIF cells expressing indicated sgRNAs (n = 4 mice in sgNT group, n = 5 mice in sg*Bcor*/*Zc3h12a* group). **e, f, k** and **l**, data represent mean ± SEM from one of three independent experiments. \**P* < 0.05, \*\*\*\**P* < 0.0001, ns, not significant, one-way ANOVA multiple-comparisons test in (**e,f**), two-way ANOVA multiple-comparisons test in (**k**), two-tailed unpaired Student’s t test in (**l**).

CD19 is a self-protein expressed by mature B cells and their progenitors, which makes CAR19T cells autoreactive by nature. The reasons why CAR19T cells cannot expand in immunocompetent mice remain unclear ^24^. However, in autoimmune diseases, autoreactive T cells expand efficiently and cause sustained tissue damages in lymphoreplete mice and human ^25^. This paradox made us wondering whether we could harness the mechanisms of autoimmunity to expand CAR19T cells in immunocompetent mice without conditioning. We screened literatures for genes whose loss-of-function in effector T cells could induce spontaneous expansion of T cells. Based on the severity of autoimmune phenotypes of mice, we ranked *Bcl2l11*/*Fas* double deficiency ^26, 27, 28^, *Ctla4*- deficiency ^29^, *Tgfbr2*-deficiency ^30^, and *Zc3h12a*-deficiency ^31^ as top candidates.

We constructed a vector expressing sgRNA and mCD19 CAR from independent promoters with a Thy1.1 marker (Fig. 1b), which could generate CAR19T cells with gene targeted by sgRNA in one step after delivering into Cas9-expressing T cells. Using this system, we produced CAR19T cells with sgRNA targeting *Bcl2l11*/*Fas*, *Ctla4*, *Tgfbr2* or *Zc3h12a*, with non-targeting (NT) sgRNA as a negative control (Fig. 1c). After transfer of these gene-targeted CD8^+^ CAR19T cells into C57BL/6 (B6) mice, only CAR19T cells with sgRNA targeting *Zc3h12a* expanded and cleared endogenous CD19^+^ B cells on day 7 post-transfer (Fig. 1d-f). However, after 4 weeks, ZC3H12A-deficient CAR19T cells disappeared and B cells started to rebound (Fig. 1d-f), a situation mimicking relapse after CAR19T cells therapy ^23^. Thus, although ZC3H12A-deficiency can boost the expansion of CAR19T cells that transiently eliminate target cells, these cells do not persist to form memory.

We next searched additional gene whose loss-of-function could increase the persistence of ZC3H12A-deficient CAR19T cells. To achieve this, we modified our vector to express two sgRNAs and mCD19 CAR simultaneously (Fig. 1g). One sgRNA was fixed to target *Zc3h12a*, while the other was a genome-wide sgRNA library targeting protein-coding genes in mouse genome (Fig. 1g) ^32^. We transferred ∼ 150 million of CAR19T cells with sgRNAs targeting *Zc3h12a* and another unknown gene into 50 mice (3 million/mouse), and harvested CD8^+^Thy1.1^+^ cells from these mice after 7 days and 3 months post-transfer for deep sequencing analysis of sgRNA enrichment (Fig. 1g).

The top hit from our screening was BCOR (Fig. 1h), a transcription repressor without known roles in T cell persistence ^33^. For validation, we transferred CAR19T cells expressing sgNT (non-targeting), sg*Zc3h12a*, or sg*Bcor* and sg*Zc3h12a* (sg*Bcor*/*Zc3h12a*) into B6 mice and monitored B cells and CAR19T cells from peripheral blood at different time points (Fig. 1i). As expected, ZC3H12A-deficient CAR19T cells contracted quickly after expansion (Fig. 1j,k). In contrast, ZC3H12A- and BCOR-double deficient CAR19T cells expanded to a similar extent as ZC3H12A-deficient CAR19T cells, but showed limited contraction (Fig. 1j,k). Consistently, there was no B cell rebound in peripheral blood of these mice after 6 months (Fig. 1l). As a control, BCOR-deficiency alone could not expand CAR19T cells (Extended Data Fig. 1d-f), indicating BCOR- deficiency must act on top of ZC3H12A-deficiency to promote the persistence of CAR19T cells. The editing of *Zc3h12a* gene was validated by immunoblot (Extended Data Fig. 1g), and the editing of *Bcor* genes was validated by DNA sequencing due to lack of good antibody for mouse BCOR protein (Extended Data Fig. 1h). Although initial editing efficiency of *Bcor* (two days post-transduction) was relatively low, *Bcor* locus in all CD8^+^Thy1.1^+^ cells isolated from mice one month post-transfer were edited (Extended Data Fig. 1i,j), indicating strong selection for cells devoid of BCOR.

Together, two rounds of screening reveal that repression of two genes in mouse genome, *Zc3h12a* and *Bcor*, generated CAR19T cells that can expand, persist and confer long-term B cell depletion in immunocompetent mice without conditioning. For simplicity and reasons described below, we named CAR19T cells devoid of ZC3H12A and BCOR as CAR19TIF(immortal-like and functional) cells, reflecting their immortal-like and functional state (see below).

### CAR19TIF cells possess almost infinite stemness but are not transformed

To test the stemness of CAR19TIF cells, we performed serial transfer experiments (Fig. 2a). During each transfer, ∼ 2 million of CAR19TIF cells from last generation of recipients were transferred into a new batch of mice (Fig. 2a). Unexpectedly, after 6 successive transfers, the percentages and cell numbers of splenic CAR19TIF cells from the 6° recipients were similar to that of 1° recipients (Fig. 2b-d). All CD19^+^ cells were eradicated in each generation of recipients (Fig. 2e), demonstrating CAR19TIF cells are functional during serial transfers. HSCs, the bona fide adult stem cells, cease to replicate and repopulate after 3 to 4 serial transfers ^34, 35^. Thus, the self-renewing capacity of CAR19TIF cells surpasses HSCs and resembles iPSCs that self-renew infinitely ^36^. Nevertheless, CAR19TIF cells could not survive in vitro (Fig. 2f), nor in NSG mice (see below), demonstrating CAR19TIF cells are not transformed.

**Fig. 2.**
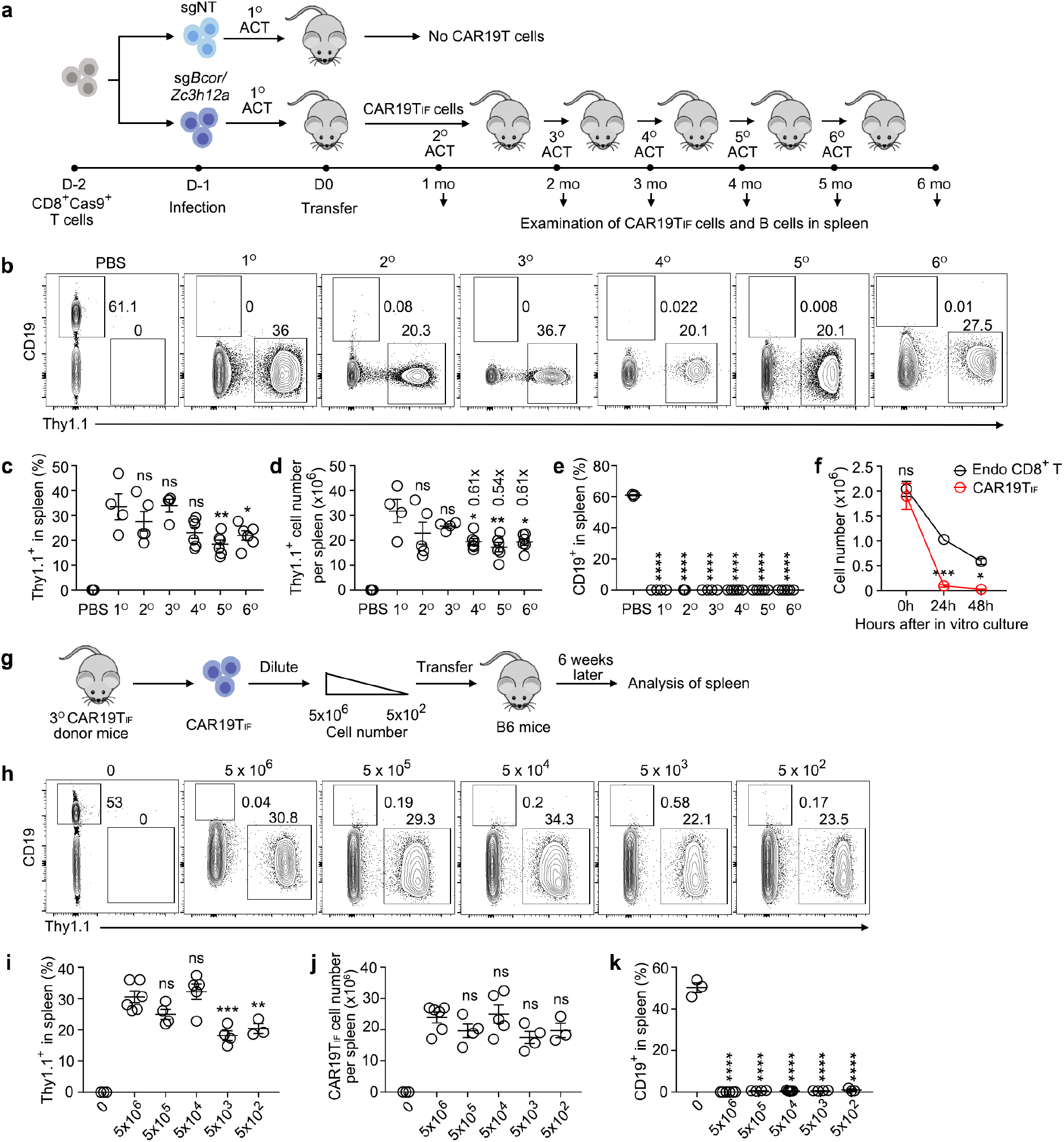
CAR19TIF cells are immortal-like and super-functional, which have a saturable niche in vivo but cannot survive in vitro. **a,** Experimental design of serial transfer of CAR19TIF cells in B6 mice. **b-e**, Representative plots (**b**) and statistical analysis (**c**-**e**) of Thy1.1^+^ CAR19TIF cells and CD19^+^ B cells from spleen 1 month post-transfer are shown (n = 3 mice in PBS group, n = 4 mice in 1° group, n = 5 mice in 2° group, n = 4 mice in 3° group, n = 6 mice in 4° group, n = 6 mice in 5° group, n = 6 mice in 6° group). **f**, Survival of CAR19TIF cells in vitro. CAR19TIF and endogenous CD8^+^ T cells isolated from mice receiving CAR19TIF cells 1 month post-transfer were cultured in vitro in T cell medium for indicated hours, and the number of live cells were counted (n = 4 replicates for each group). **g**, Experimental design of transferring different numbers of CAR19TIF cells into B6 mice. **h**-**k**, Representative plots (**h**) and statistical analysis (**i**-**k**) of Thy1.1^+^ CAR19TIF cells and CD19^+^ B cells from spleen 1 month post-transfer are shown (n = 3 mice in PBS group, n = 6 mice in 5×10^6^ group, n = 4 mice in 5×10^5^ group, n = 5 mice in 5×10^4^ group, n = 4 mice in 5×10^3^ group, n = 3 mice in 5×10^2^ group). **c**-**f** and **i**-**k**, data represent mean ± SEM from one of three independent experiments. \**P* < 0.05, \*\**P* < 0.01, \*\*\**P* < 0.001, \*\*\*\**P* < 0.0001, ns, not significant, one-way ANOVA multiple-comparisons test in (**c**-**e** and **i**-**k**), two-way ANOVA multiple-comparisons test in (**f**).

Six months after transfer of CAR19TIF cells, all recipient mice looked healthy (Extended Data Fig. 2a), and had normal body weights (Extended Data Fig. 2b). Mice transferred with CAR19TIF cells did not show sign of cell overgrowth or inflammatory response in spleen (Extended Data Fig. 2c). Endogenous CD8^+^ T cell numbers were slightly increased in mice transferred with CAR19TIF cells (Extended Data Fig. 2d,e), likely due to absence of B cells. Most endogenous CD8^+^ T cells maintained a naive phenotype in the presence of CAR19TIF cells (Extended Data Fig. 2f,g). These data demonstrate that, although CAR19TIF cells persist in vivo in large quantities (∼ total endogenous CD8^+^ T cell number) (Extended Data Fig. 4d-f), they do not cause obvious side-effects.

To test whether the stemness of CAR19TIF cells is conferred by a small population of cells with less proliferation, we coupled serial transfers with CFSE labelling to track cell division (Extended Data Fig. 3a). Theoretically, CAR19TIF cells with high level of CFSE are expected to proliferate less and likely retain stemness. We found that, in each generation of transfer, CAR19TIF cells with lowest level of CFSE were still able to proliferate extensively and repopulate next generation of recipients (Extended Data Fig. 3b,c). These data demonstrate that CAR19TIF cells that have experienced extensive proliferation in the last generation of hosts still retain stemness to repopulate naive hosts. Analysis of CAR19TIF cells before and after transfer showed that these cells underwent reversible phenotypic changes. Before transfer, CAR19TIF cells from spleen were CD62L^+^CD69^-^ and smaller than that of endogenous CD8^+^ T cells (Extended Data Fig. 3d-i). After transfer into new recipients, CAR19TIF cells were activated rapidly with increase of cell size, upregulation of CD69 and partial downregulation of CD62L (Extended Data Fig. 3g-i). This activation process could be mimicked by stimulation of CAR19TIF cells by B cells in vitro (Extended Data Fig. 3j). Four weeks after transfer, all these phenotypic changes were reversed to pre-transfer level (Extended Data Fig. 3g-i). However, CAR19TIF cells showed limited contraction after extensive expansion (Extended Data Fig. 3d-f).

To exclude off-target effects of sgRNAs targeting *Bcor* and/or *Zc3h12a*, we used a different set of sgRNAs to induce CAR19TIF cells. CAR19TIF cells could be induced, maintained and serially transferred in B6 mice with this different set of sgRNAs (data not shown), demonstrating the induction of CAR19TIF cells is not due to off-target effects of sgRNAs. Together, these data demonstrate that CAR19TIF cells possess almost infinite self-renewing capacity like iPSCs but retain functionality of mature T cells.

### CAR19TIF cells are super-functional and have a saturable niche in vivo

We transferred different numbers of CAR19TIF cells into B6 mice, from 5, 000, 000 to 500 cells, and analyzed cells recovered from spleen after 6 weeks (Fig. 2g). Surprisingly, in spite of 10,000-fold difference of cell input, CAR19TIF cells showed similar percentages and cell numbers among different groups (Fig. 2h-j), demonstrating CAR19TIF cells have a saturable niche in vivo. As few as 500 of CAR19TIF cells were sufficient to eliminate all the hundreds of millions of endogenous B cells in recipient mice in the absence of any conditioning (Fig. 2k), and reconstitute their own compartment (Fig. 2h-j), demonstrating superior functionality of CAR19TIF cells.

We analyzed the distribution of CAR19TIF cells in mice (Extended Data Fig. 4a,b). No CD19^+^ cells could be detected in mice with CAR19TIF cells (Extended Data Fig. 4c), and CAR19TIF cells were present in all organs examined with underrepresentation in lymph nodes (LNs) and relative enrichment in bone marrow (BM) (Extended Data Fig. 4b-g). CAR19TIF cells in BM expressed higher levels of CD25 and PD-1, and were slightly bigger than those in spleen (Extended Data Fig. 4h,i). About ∼ 1% of CAR19TIF cells from spleen were in cell cycle (Ki67^+^), while 7-fold more such cells were found in BM (Extended Data Fig. 4j,k). About a quarter of CAR19TIF cells from BM were CD69^+^, a marker of recent activation or tissue residency ^37, 38^, while only 4% of such cells were present in spleen (Extended Data Fig. 4l,m). The expression of stemness marker CD62L and CXCR5 on CAR19TIF cells was comparable between spleen and BM (Extended Data Fig. 4h,i,n,o). Thus, although no target cells could be detected, CAR19TIF cells showed broad distribution in mice with relative enrichment in BM.

To explore the potential role of the undetectable CD19^+^ progenitor cells in the maintenance of CAR19TIF cells, we used *Rag1*^-/-^ mice and NSG mice as recipients. *Rag1*^-/-^ mice barely had CD19^+^ mature B cell in spleen but had ∼ 3% of CD19^+^ progenitor cells in BM (Extended Data Fig. 4p,q), while NSG mice were completely devoid of CD19^+^ cells (Extended Data Fig. 4r,s). CAR19TIF cells could expand, kill CD19^+^ cells and persist in *Rag1*^-/-^ but not NSG mice (Extended Data Fig. 4t-w), suggesting the expansion and persistence of CAR19TIF cells require the presence of CD19^+^ target cells.

Taken together, these data demonstrate that CAR19TIF cells are super-functional and have a saturable niche in vivo, and they cannot expand nor persist in mice devoid of CD19^+^ cells.

### CAR19TIF cells exhibit hybrid features of effector, memory and precursor exhausted T cells

In mechanistic studies, we could not obtain appropriate controls for CAR19TIF cells due to their striking persistence and stemness. As we showed above, wild-type and BCOR- deficient CAR19T cells did not expand in B6 mice (Extended Data Fig. 1a-f), while ZC3H12A-deficient CAR19T cells contracted within 2 weeks (Fig. 1j,k). Thus, we used endogenous CD8^+^ T cells as a control for phenotypic analysis since these cells were not affected by CAR19TIF cells (Extended Data Fig. 2f,g). Two months post-transfer, the majority of endogenous splenic CD8^+^ T cells from mice with CAR19TIF cells were CD62L^hi^CD44^lo^ naïve T (TN) cells, and ∼ 10% were CD62L^hi^CD44^hi^ TCM cells (Fig. 3a,b). About 95% of CAR19TIF cells in spleen were CD62L^hi^CD44^hi^ (Fig. 3a,b), a phenotype of TCM cells. The expression levels of CD62L, TCF1, CD25, CD127 and CD122 were comparable between CAR19TIF cells and endogenous CD8^+^ T cells (Fig. 3c,d and Extended Data Fig. 5), again suggesting TN or TCM cell phenotype of CAR19TIF cells. The increased expression of ICOS was expected (Extended Data Fig. 5a,b), since it is negatively regulated by ZC3H12A ^31^. CD27 and CD28 were also increased on CAR19TIF cells (Extended Data Fig. 5c,d). CAR19TIF cells expressed high levels of PD-1 and CXCR5 (Fig. 3c,d), markers of follicular helper T cells or TPEX cells ^12, 13, 15^. The expression of LAG-3 and TIM-3 was only minimally increased on CAR19TIF cells compared with endogenous CD8^+^ T cells (Extended Data Fig. 5a,b), suggesting these cells are not terminally exhausted, which agrees with expansion and killing capacity of these cells (Fig. 2). CAR19TIF cells also expressed c-Kit, a marker of HSCs (Fig. 3c,d), but had low level of CD150 (Extended Data Fig. 5c,d), a marker of TSCM cells ^6, 7^. In vitro stimulation with phorbol 12-myristate 13-acetate (PMA) and ionomycin showed that CAR19TIF cells produced IL-2 and IFNγ, similar to that of TCM cells (Fig. 3e,f). In vitro killing assay revealed that CAR19TIF cells could kill B cells directly without pre-expansion (Fig. 3g), demonstrating these cells are cytotoxic.

**Fig. 3.**
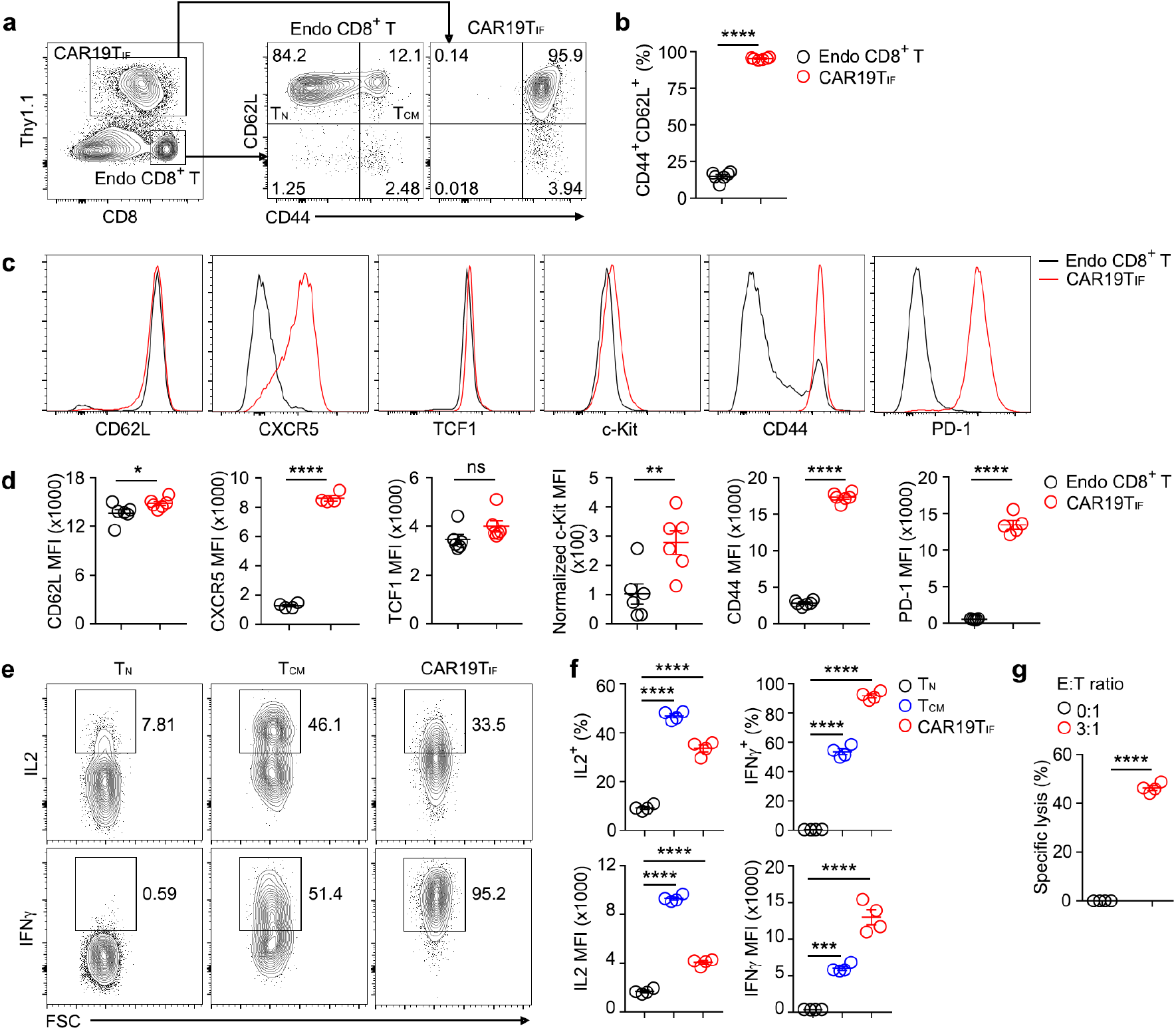
CAR19TIF cells exhibit hybrid features of effector, memory and precursor exhausted T cells. **a,** Flow cytometry gating of endogenous CD8^+^ T cells and CAR19TIF cells from spleen of mice 2 months post-transfer. **b**, Statistical analysis of the percentages of CD44^+^CD62L^+^ cells among endogenous CD8^+^ T cells and Thy1.1^+^ CAR19TIF cells (n = 6 mice for each group). **c,d**, Flow cytometry analysis of the expression of indicated protein on endogenous CD8^+^ T cells and Thy1.1^+^ CAR19TIF cells 2 months post-transfer. Representative plots (**c**) and statistical analysis of mean florescence intensity (MFI) (**d**) are shown (n = 4 mice for CXCR5 panel, n = 6 mice for other groups). **e,f**, Cells were stimulated with 50 ng/ml phorbol 12-myristate 13-acetate (PMA) and 1 µM ionomycin for 4 hours in the presence of GolgiStop, expression of IL2 and IFNγ were examined by intracellular staining. Representative plots (**e**) and statistical analysis (**f**) are shown (n = 4 mice for each group). **g**, Purified B cells were co-cultured with CAR19TIF cells isolated from spleen of 2° mice 2 months post-transfer or medium control at indicated ratio for 24 hours. After killing, live B cells were examined by flow cytometry and quantified (n = 4 replicates for each group). **b, d, f** and **g**, data represent mean ± SEM from one of three independent experiments. *p < 0.05, **p < 0.01, ****p < 0.0001, ns, not significant, two-tailed unpaired Student’s t test in (**b,d,g**), one-way ANOVA multiple-comparisons test in (**f**).

These data demonstrate that CAR19TIF cells exhibit a unique phenotype and do not belong to known T cell subset, and the simple and unique CD44^hi^CD62L^hi^PD-1^hi^ phenotype distinguishes CAR19TIF cells from other T cell subsets (see **Discussion**).

### ZC3H12A- and BCOR-deficiency synergistically reprogram CAR19TIF cells

To explore the mechanisms of CAR19TIF cells reprograming, we first performed bulk RNA-seq analysis of cells isolated from spleen. Principle component analysis (PCA) showed that, on day 10 post-transfer when ZC3H12A-deficient CAR19T cells were contracting while ZC3H12A- and BCOR-double deficient CAR19TIF cells persisted (Fig. 1k), the transcriptome of ZC3H12A-deficient CAR19T cells and CAR19TIF cells already segregated from each other (Fig. 4a). Compared with ZC3H12A-deficiency alone, CAR19TIF cells had 1,183 genes downregulated and 239 genes upregulated (Extended Data Fig. 6a). Pathway analysis showed that inflammation-associated modules were repressed in CAR19TIF cells while signaling associated with pluripotency, stemness and Wnt pathway were upregulated (Extended Data Fig. 6b,c), indicating BCOR-deficiency represses inflammation but induces stemness.

**Fig. 4.**
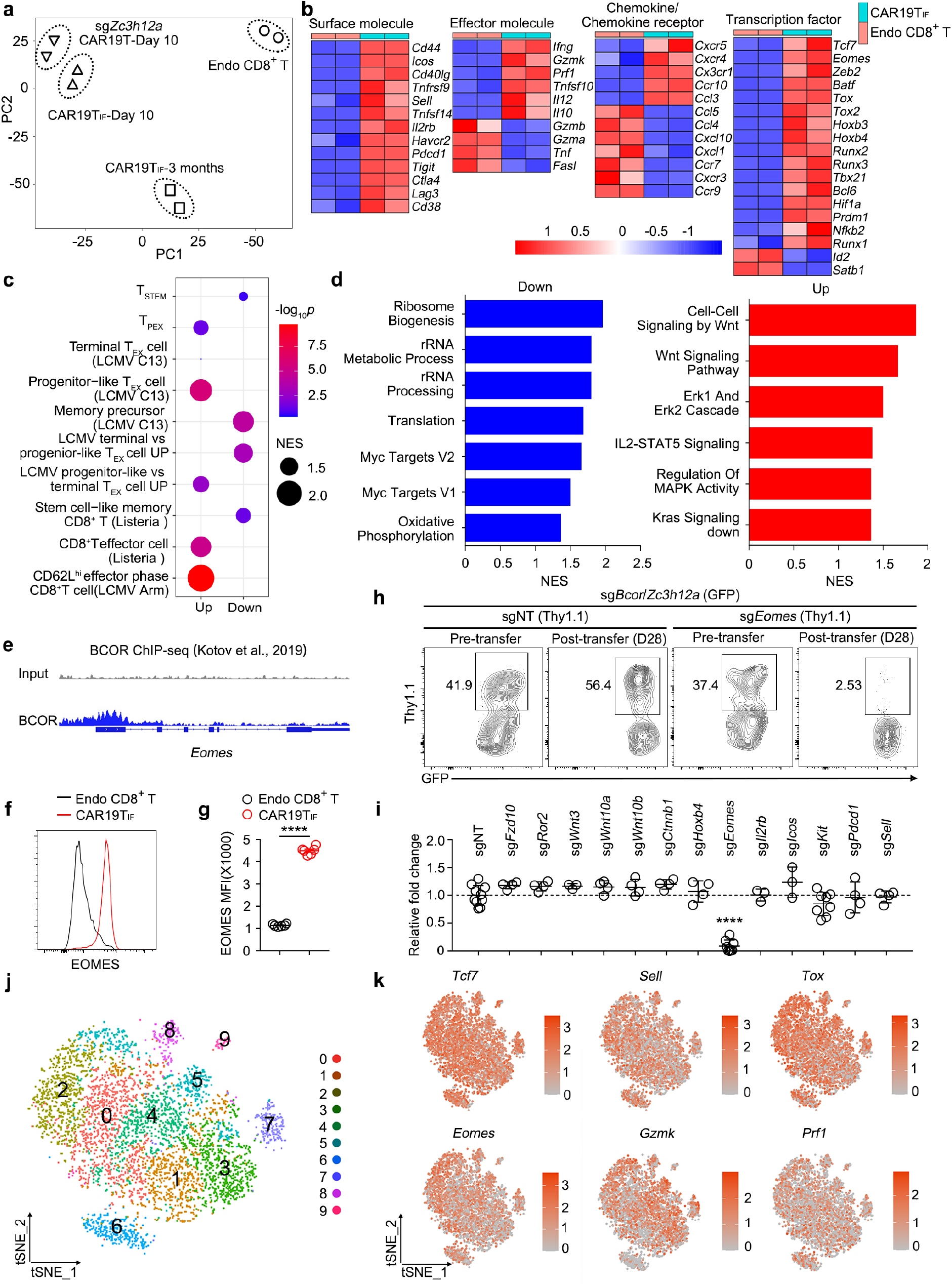
ZC3H12A- and BCOR-deficiency synergistically reprogram CAR19TIF cells. **a,** PCA of ZC3H12A-deficient CAR19T cells 10 days post-transfer, CAR19TIF cells 10 days post-transfer, CAR19TIF cells 3 months post-transfer and endogenous CD8^+^ T cells 3 months post-transfer. **b**, Heatmaps showing the expression of selected genes in CAR19TIF cells and endogenous CD8^+^ T cells 3 months post-transfer. **c**, Similarity of CAR19TIF cells (3 months post-transfer) with other T cell subsets from published studies. **d**, Pathway analysis of upregulated and downregulated genes in CAR19TIF cells (3 months post-transfer) compared with endogenous CD8^+^ T cells. **e**, ChIP-seq data of BCOR binding to *Eomes* promoter region. **f,g**, Flow cytometry analysis of EOMES expression in endogenous CD8^+^ T cells and Thy1.1^+^ CAR19TIF cells (one month post-transfer). Representative plots (**f**) and statistical analysis of mean florescence intensity (MFI) (**g**) are shown (n = 6 mice for each group). **h,i**, CD8^+^Cas9^+^ T cells were activated, co-infected with retrovirus expressing CAR19 with sgRNA targeting *Bcor*/*Zc3h12a* (GFP^+^) and retrovirus expressing CAR19 with sgRNA targeting another indicated gene. One million of CD8^+^GFP^+^ CAR19T cells were transferred into B6 mice. After 4 weeks, the ratio of GFP^+^Thy1.1^+^ cells among total GFP^+^ cells from spleen were examined. Data were normalized to pre-transfer. Representative plots (**h**) and statistical analysis (**i**) are shown (n = 10 mice in sgNT group, n = 4 mice in sg*Fzd10* group, n = 4 mice in sg*Ror2* group, n = 3 mice in *Wnt3* group, n = 4 mice in sg*Wnt10a* group, n = 4 mice in sg*Wnt10b* group, n = 4 mice in sg*Ctnnb1* group, n = 4 mice in sg*Hoxb4* group, n = 8 mice in sg*Eomes* group, n = 3 mice in sg*IL2rb* group, n = 3 mice in sg*Icos* group, n = 8 mice in sg*kit* group, n = 4 mice in sg*Pdcd1* group, n = 4 mice in sg*Sell* group). **j**, Unsupervised clustering of CAR19TIF cells by scRNA-seq. **k**, The expression of indicated genes in CAR19TIF cells at single cell level. **g** and **i**, data represent mean ± SEM from one of two independent experiments. \*\*\*\**P* < 0.0001, two-tailed unpaired Student’s t test.

Three months post-transfer, when ZC3H12A-deficient CAR19T cells disappeared and CAR19TIF cells in spleen entered a quiescent state, the transcriptome of CAR19TIF cells was highly different from that of endogenous CD8^+^ T cells, with 1,079 genes downregulated and 1,479 genes upregulated (Extended Data Fig. 6d). Several co-stimulatory molecules were upregulated in CAR19TIF cells, including ICOS, 4-1BB and TIGIT (Fig. 4b). GZMB and TNF were downregulated while IFNγ and GZMK were upregulated (Fig. 4b), consistent with cytotoxic function of these cells (Fig. 3g). The lymph node homing receptor CCR7 was downregulated in CAR19TIF cells (Fig. 4b), explaining low abundance of these cells in lymph node (Extended Data Fig. 4b). The expression of several inflammatory chemokine receptors including CXCR4 and CX3CR1 were increased in CAR19TIF cells (Fig. 4b), consistent with their whole-body distribution (Extended Data Fig. 4b). For transcription factors (TFs), the expression of TCF1, TOX, TOX2, BATF, ZEB2, BLIMP1, BCL6 and EOMES were increased in CAR19TIF cells compared with endogenous CD8^+^ T cells, while ID2 and SATB1 were downregulated (Fig. 4b). In addition, CAR19TIF cells also expressed HOXB3 and HOXB4 (Fig. 4b), HSCs-specific TFs that usually do not present in T cells ^39^. Comparing the transcriptome of CAR19TIF cells with RNA-seq data of known T cell subsets showed that CAR19TIF cells exhibited mixed feature of stem-like T cells, TPEX cells and effector T cells (Fig. 4c and Extended Data Fig. 6e) ^40, 41, 42, 43^. Pathway analysis revealed that Wnt signaling was highly enriched while ribosome biogenesis, translation, MYC targets and oxidative phosphorylation (OXPHOS) were repressed in CAR19TIF cells (Fig. 4d), further supporting stem cell feature and quiescence of CAR19TIF cells. It is noticeable that the transcriptional profiles of CAR19TIF cells between 10 days and 3 months after transfer differed significantly (Fig. 4a and Extended Data Fig. 6f), indicating a differentiation process after clearance of target cells. During this process, inflammation-, cell cycle- and metabolism-associated programs were downregulated while stemness-associated modules were further enhanced (Extended Data Fig. 6g-i).

Published ChIP-seq data showed that *Eomes* was a direct target of BCOR (Fig. 4e) ^44^. We validated that EOMES protein was highly upregulated in CAR19TIF cells (Fig. 4f,g). To dissect the reprogramming mechanism of CAR19TIF cell, we performed rescue experiments to interrogate the role of upregulated genes in CAR19TIF cells, including multiple Wnt pathway components (*Fzd10*, *Ror2*, *Wnt3*, *Wnt10a*, *Wnt10b* and *Ctnnb1*), as well as *Hoxb4*, *Eomes*, *Il2rb*, *Icos*, *Kit*, *Pdcd1* and *Sell*. Among these genes, *Eomes* was required for the expansion and/or persistence of CAR19TIF cells (Fig. 4h,i).

To explore whether the hybrid feature of CAR19TIF cells is due to population heterogeneity, we performed single cell RNA sequencing (scRNA-seq) analysis of CAR19TIF cells from spleen (Extended Data Fig. 7a). Although CAR19TIF cells were partitioned into 10 “clusters” using unsupervised clustering (Fig. 4j and Extended Data Fig. 7b), the differences of gene expression among different “clusters” were minimal (Extended Data Fig. 7b). Importantly, the expression of key functional genes was highly similar among “clusters”, including *Tcf7*, *Sell*, *Tox*, *Eomes*, *Gzmk*, *Prf1* and others (Fig. 4k and Extended Data Fig. 7c). In addition, topic modeling analysis showed that stemness-associated topic (topic5) scattered among all “clusters” (Extended Data Fig. 7d,e) ^45^, indicating the stemness of CAR19TIF cells does not reside in certain subset. The expression of stemness and effector function genes in the same cell demonstrates that the hybrid feature of CAR19TIF cells is at single cell level.

Taken together, these data suggest ZC3H12A- and BCOR-deficiency synergistically induce a unique program in CAR19TIF cells that confers these cells expansion, stemness and quiescence (see Fig. 8k and **Discussion**).

### CAR19TIF cells enter a metabolically dormant state after elimination of target cells

Metabolism plays important roles in T cell fate decision ^46^. On day 7 post-transfer, the expanding CAR19TIF cells were much bigger than that of endogenous CD8^+^ T cells (Extended Data Fig. 8a). However, after 4 weeks when CAR19TIF cells returned back to quiescence, splenic CAR19TIF cells were even smaller than TN cells (Extended Data Fig. 8a). This dynamic change of cell size of CAR19TIF cells was maintained during serial transfers (Extended Data Fig. 8b), demonstrating smaller size in the absence of target cells is an intrinsic property of CAR19TIF cells. Proteins, particularly ribosomes, constitute the major part of cell mass ^47^. We found ribosome biogenesis was repressed in splenic CAR19TIF cells compared with endogenous CD8^+^ T cells (Extended Data Fig. 8c). Indeed, the expression of most ribosome components was significantly downregulated in CAR19TIF cells (Extended Data Fig. 8d). MYC, the master transcription factor for ribosome biogenesis and metabolism ^47^, was reduced in CAR19TIF cells compared with TN cells (Extended Data Fig. 8e,f).

Engaging mitochondrial metabolism is metabolic feature of long-lived T cells ^46^. However, our RNA-seq data showed that genes related to OXPHOS were downregulated in CAR19TIF cells compared with endogenous CD8^+^ T cells (Extended Data Fig. 8g). Seahorse experiments showed that oxygen consumption rate (OCR) and extracellular acidification rate (ECAR) were both reduced in CAR19TIF cells (Extended Data Fig. 8h,i). Consistently, glucose uptake was markedly reduced in CAR19TIF cells compared with endogenous CD8^+^ T cells (Extended Data Fig. 8j,k), which explains low metabolic activity of CAR19TIF cells.

Untargeted metabolome analysis showed that ∼ 319 metabolites were increased while ∼309 metabolites were decreased in CAR19TIF cells (Extended Data Fig. 8l), which metabolically segregated CAR19TIF cells from endogenous CD8^+^ T cells (Extended Data Fig. 8m). Adenosine was one of the highly increased metabolites in CAR19TIF cells (Extended Data Fig. 8l,n). Adenosine is an immunosuppressive metabolite produced by nucleosidases CD39 and CD73 ^48^, which has been shown to inhibit glucose uptake and metabolism of T cells ^49^. We found that both CD73 (encoded by *Nt5e*), the enzyme producing adenosine, and ADORA2A, the receptor for adenosine, were upregulated in CAR19TIF cells (Extended Data Fig. 8o-r). Treatment with ADORA2A antagonist ZM241385 partially increased glucose uptake of CAR19TIF cells (Extended Data Fig. 8s), suggesting autocrine production of adenosine is one of the mechanisms how CAR19TIF cells maintain metabolic quiescence.

These data demonstrate that, although CAR19TIF cells showed little contraction after expansion, they enter a dormant state with minimal metabolic activities.

### CAR19TIF cells mediate long-term repression of tumors

Poor expansion and/or persistence of adoptively transferred T cells is a barrier for durable ACT ^4, 5^. In our system, wild-type and BCOR-deficient CAR19T cells did not expand in immunocompetent mice (Extended Data Fig. 1a-f), and ZC3H12A-deficient CAR19T cells could not persist more than 2 weeks (Fig. 1k). However, CAR19TIF cells persisted in mice and these mice never showed B cell rebound during the observation period (Fig. 1k,l). Consistently, after transfer of congenically marked splenocytes (CD45.1^+^) into control B6 mice (CD45.2^+^) or B6 mice previously received CAR19TIF cells (Fig. 5a), CD19^+^ B cells could only be detected in control mice but not mice with CAR19TIF cells, while engraftment of CD19^-^ non-B cells were detected in both groups of mice (Fig. 5b,c), demonstrating that CAR19TIF cells keep monitoring and killing endogenous and exogenous CD19^+^ cells.

**Fig. 5.**
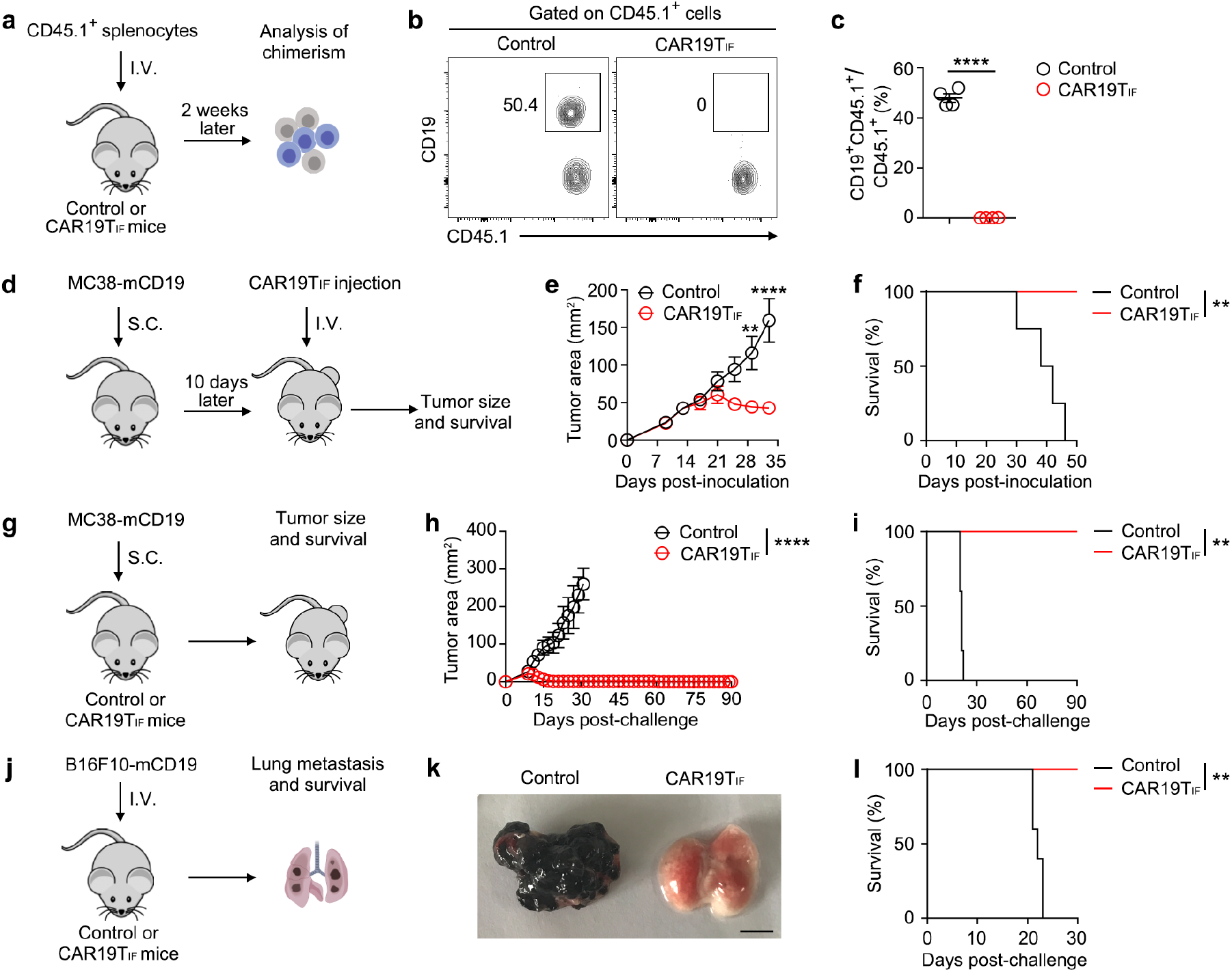
CAR19TIF cells mediate long-term repression of tumors. **a,** Experimental design. Ten million of splenocytes from CD45.1 mouse were injected into control B6 mice or B6 mice with CAR19TIF cells (male) via tail vein. Chimerism of splenocytes was analyzed by flow cytometry 2 weeks post-transfer. **b,c**, Representative plots (**b**) and statistical analysis (**c**) of CD45.1^+^CD19^+^ B cells among total CD45.1^+^ cells in spleen are shown (n = 4 mice for each group). **d**, Experimental design. Male B6 mice were subcutaneously inoculated with MC38-mCD19 cells (0.2 million). Ten days later, CAR19TIF (3 million per mice) from 2° donor mice or PBS were transferred into tumor bearing mice. Tumor growth and mice survival were monitored. **e,f**, Tumor growth (**e**) and survival (**f**) of mice with indicated treatment (n = 4 mice for each group). **g**, Experimental design. MC38-mCD19 cells (0.5 million) were subcutaneously inoculated in control B6 mice or mice with CAR19TIF (male). Tumor growth and mice survival were monitored. **h,i**, Tumor growth (**h**) and survival (**i**) of mice with indicated treatment (n = 5 mice in control group, n = 10 mice in CAR19TIF group). **j**, Experimental design. B16F10-mCD19 cells (0.1 million) were injected into control B6 mice or B6 mice with CAR19TIF cells (male) via tail vein. Tumor growth and mice survival were monitored. **k,l**, Representative image of tumor burden in lung 3 weeks post-transfer (**k**) and mice survival (**l**) are shown (n = 5 mice in control group, n = 6 mice in CAR19TIF group). Bar is 0.5 cm. **c, e** and **h**, data represent mean ± SEM from one of two independent experiments. \*\**P* < 0.01, \*\*\*\**P* < 0.0001, two-tailed unpaired Student’s t test in (**c**), two-way ANOVA multiple-comparisons test in (**e,h**), log-rank (Mantel-Cox) test in (**f,i,l**).

We then tested CAR19TIF cells in primary and memory response against tumors. We did not use the non-expanding BCOR-deficient CAR19T cells (Extended Data Fig. 1d-f) and non-persistent ZC3H12A-deficient CAR19T cells (Fig. 1k) as controls, due to their obvious defects in long-term elimination of endogenous CD19^+^ cells. Mice mock transferred with PBS were used as controls. In primary response (Fig. 5d), CAR19TIF cells repressed the growth of MC38 tumor cells expressing mCD19 (MC38-mCD19) and extended mice survival (Fig. 5e,f). To test CAR19TIF cells in memory protection, we inoculated MC38-mCD19 subcutaneously into control mice and mice previously received CAR19TIF cells (Fig. 5g). Although MC38-mCD19 tumor grew rapidly in control B6 mice, mice with CAR19TIF cells showed transient tumor growth followed by regression (Fig. 5h), which prolonged survival of mice with CAR19TIF cells (Fig. 5i). Metastasis is common in tumor relapse, which is a major cause of death. B16F10 melanoma cells expressing mCD19 (B16F10-mCD19) were intravenously injected into mice via tail vein, which is a commonly used model of lung metastasis (Fig. 5j). We found B16F10-mCD19 cells quickly colonized lungs of control mice, resulting in rapid death of mice (Fig. 5k,l). In contrast, B16F10-mCD19 cells could not efficiently colonize lungs of mice with CAR19TIF cells (Fig. 5k), and these mice lived longer (Fig. 5l). Collectively, these data demonstrate that CAR19TIF cells mediate long-term protection against tumors.

### Induction of TIF cells with CARs targeting solid tumor antigens

We next investigated whether TIF cells could be induced by CARs targeting other antigens. We first tested a CAR targeting human EGFR (derived from the Cetuximab antibody) (Fig. 6a) ^50^. EGFR CAR-T cells expressing sgNT, sg*Bcor*, sg*Zc3h12a* or sg*Bcor*/*Zc3h12a* were transferred into B6 mice, and the expansion and persistence of CAR-T cells were monitored (Fig. 6b). Serial bleeding showed that only EGFR CAR-T cells expressing sgRNAs targeting both *Bcor* and *Zc3h12a* were detected in peripheral blood, which peaked at 4 weeks post-transfer and persisted for 24 weeks at least (Fig. 6c). Consistently, only when both BCOR and ZC3H12A were depleted, were EGFR CAR-T cells detected in spleen (Fig. 6d-f). These data indicate that, unlike in CAR19T cells where ZC3H12A-deficiency alone promoted CAR-T cells expansion (Fig. 1d-f), ZC3H12A- deficiency alone cannot promote EGFR CAR-T cells expansion while BCOR- and ZC3H12A-double deficiency can enhance the expansion and persistence of EGFR CAR-T cells, named as EGFRTIF cells for their resemblance of CAR19TIF cells (see below). Flow cytometry analysis of splenic EGFRTIF cells 4 weeks post-transfer revealed a CD44^hi^CD62L^hi^ central memory-like phenotype (Extended Data Fig. 9a,b). EGFRTIF cells expressed CXCR5, TCF1, PD-1 but not TIM-3 (Extended Data Fig. 9c), typical features of CAR19TIF (Fig. 3 and Extended Data Fig. 5). Unlike CAR19TIF cells, which were smaller than endogenous CD8^+^ T cells (Extended Data Fig. 8), EGFRTIF cells had similar cell size with endogenous CD8^+^ T cells (Extended Data Fig. 9c), suggesting EGFRTIF cells may not be as quiescent as CAR19TIF cells, likely due to tonic signaling of this EGFR CAR ^50^. EGFRTIF cells potently produced IFNγ upon ex vivo stimulation (Extended Data Fig. 9d-f), indicating these cells were not exhausted.

**Fig. 6.**
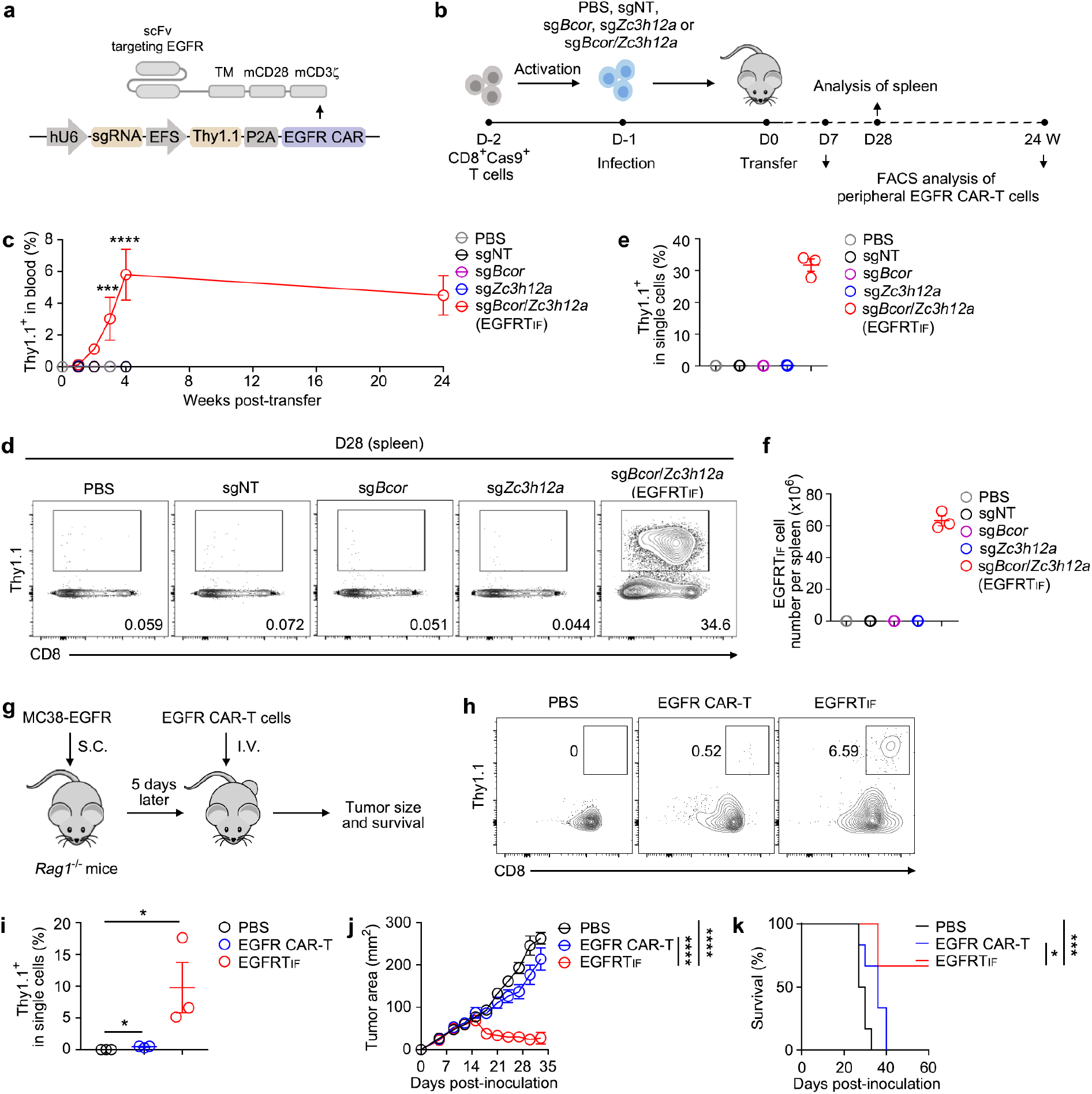
Induction of EGFRTIF cells. **a,** The composition of anti-EGFR CAR. **b**, Experimental design for induction and analysis of EGFRTIF cells. **c**, Statistical analysis of EGFRTIF cells among single live cells from peripheral blood at indicated time (n = 3 mice for each group). **d**-**f**, Representative plots (**d**) and statistical analysis (**d,f**) of EGFRTIF cells among single live cells from spleen 28 days post-transfer are shown (n = 3 mice for each group). **g**, Experimental design. *Rag1*^-/-^ mice were subcutaneously inoculated with MC38-EGFR tumor cells (0.5 million). Five days later, PBS, one million of control (sgNT) EGFR CAR-T cells or EGFRTIF cells were transferred into tumor bearing mice. Tumor growth and mice survival were monitored. **h,i**, Representative plots (**h**) and statistical analysis (**i**) of EGFR CAR-T cells among single live cells from blood are shown (n = 3 mice for each group). **j,k**, Tumor growth (**j**) and mice survival (**k**) were monitored (n = 6 mice for each group). **c, e, f** and **i**-**k**, data represent mean ± SEM. Data are representative of three independent experiments. \**P* < 0.05, \*\*\**P* < 0.001, \*\*\*\**P* < 0.0001, two-way ANOVA multiple-comparisons test in (**c,j**), one-way ANOVA multiple-comparisons test in (**i**), log-rank (Mantel-Cox) test in (**k**).

To test the stemness of EGFRTIF cells, we performed serial transfer experiments in B6 and NSG mice (Extended Data Fig. 9g). EGFRTIF cells could be serially transferred in B6 or NSG mice for at least 3 generations without outgrowth (Extended Data Fig. 9h-m), and they could not survive in vitro (Extended Data Fig. 9n). These data demonstrate that EGFRTIF cells possess superior stemness but are not transformed.

We then investigated the anti-tumor effect of EGFRTIF cells. We established MC38-EGFR tumor cells, which initially grew but were quickly rejected in B6 mice (data not shown), likely due to immunogenicity of human EGFR in mice. So, we injected MC38-EGFR cells in *Rag1*^-/-^ mice, followed by transfer of PBS, control EGFR CAR-T cells (sgNT) or EGFRTIF cells (Fig. 6g). BCOR- or ZC3H12A-single knockout EGFR CAR-T cells were not included due to their inability to expand in vivo (Fig. 6c-f). Robust expansion of EGFRTIF cells but not control EGFR CAR-T cells were detected in blood of tumor-bearing mice (Fig. 6h,i), which was associated with improved tumor control and mice survival (Fig. 6j,k).

To further test the generality of TIF program, we tested another clinically used CAR targeting glycolipid GD2 (derived from the 14G2A monoclonal antibody) (Extended Data Fig. 10a) ^51^. Due to unique framework regions (FRs) of the 14G2A antibody, this GD2 CAR showed constitutive tonic signaling when CD28 was used as the costimulation domain, resulting in CAR-T cells exhaustion and poor expansion in vivo ^51^. We transferred GD2 (14G2A) CAR-T cells expressing sgNT, sg*Bcor*, sg*Zc3h12a* or sg*Bcor*/*Zc3h12a* into B6 mice and monitored CAR-T cells (Extended Data Fig. 10b). Only when both BCOR and ZC3H12A were depleted, GD2 (14G2A) CAR-T cells expansion and persistence were observed (named as GD2TIF) (Extended Data Fig. 10c-f). GD2TIF cells exhibited almost identical phenotypes with CAR19TIF and EGFRTIF cells (Extended Data Figs. 10g- m and 9, also see Fig. 3). GD2TIF cells could be serially transferred in B6 or NSG mice for at least 3 generations without outgrowth (Extended Data Fig. 10n-t), which also could not survive in vitro (Extended Data Fig. 10u), demonstrating stemness and safety of these cells.

### Induction of GD2TIF cells requires CAR tonic signaling

The induction and maintenance of CAR19TIF cells depend on CD19^+^ target cells (Extended Data Fig. 4), while EGFR and GD2 CAR-T cells may not have large number of target cells expressing high level of cognate antigen in mice (hEGFR and GD2, respectively). We speculated that the induction of EGFRTIF and GD2TIF cells might be due to strong tonic signaling of these two CARs ^50, 51^. We thus chose another GD2 CAR (derived from the K666 monoclonal antibody), which was reported to have little tonic signaling (Fig. 7a) ^52, 53^. Depletion of BCOR and ZC3H12A only minimally expanded GD2 (K666) CAR-T cells, in contrast to robust expansion of GD2 (14G2A) CAR-T cells (GD2TIF) under the same condition (Fig. 7b-d). These data suggest that, in the absence of large amount of target cells like normal B cells for CD19 CAR, the induction of TIF program may dependent on tonic signaling of CAR. To test this hypothesis, we tried another CAR with little tonic signaling: the HER2 CAR derived from the 4D5 monoclonal antibody (Fig. 7e) ^51, 54^. Indeed, depletion of BCOR and ZC3H12A only minimally expanded HER2 CAR-T cells (Fig. 7f-h).

**Fig. 7.**
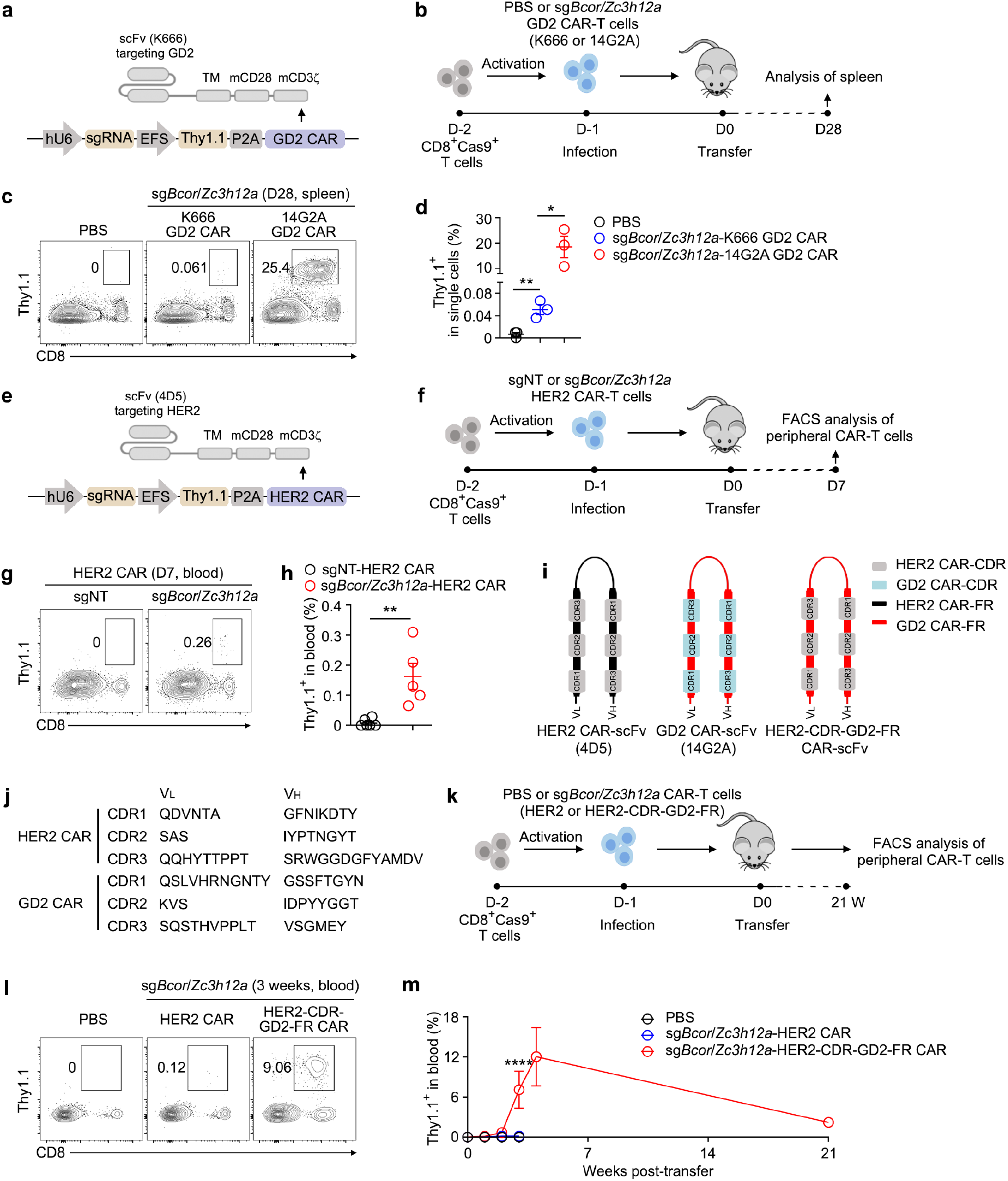
Induction of GD2TIF cells requires CAR tonic signaling. **a,** The composition of anti-GD2 CAR with scFv derived from the K666 monoclonal antibody. **b**, Experimental design. **c,d**, Representative plots (**c**) and statistical analysis (**d**) of GD2(K666) CAR-T cells and GD2(14G2A) CAR-T (GD2TIF) cells among single live cells in spleen 28 days post-transfer are shown (n = 3 mice for each group). **e**, The composition of anti-HER2 CAR with scFv derived from the 4D5 monoclonal antibody. **f**, Experimental design. **g,h**, Representative plots (**g**) and statistical analysis (**h**) of HER2 CAR-T cells among single live cells from blood 7 days post-transfer are shown (n = 5 mice for each group). **i**, scFv of HER2 CAR (4D5), GD2 CAR (14G2A) and a chimeric scFv consisting with the complementary determining regions (CDRs) of HER2 CAR and the framework regions (FRs) of the GD2 CAR (HER2-CDR-GD2-FR CAR). **j**, The CDR sequences of HER2 CAR and GD2 CAR (14G2A). **k**, Experimental design. **l,m**, Representative plots (**l**) and statistical analysis (**m**) of HER2-CDR-GD2-FR CAR CAR-T cells among single live cells from blood at indicated time are shown (n = 3 mice for each group). **d, h** and **m**, data represent mean ± SEM. Data are representative of three independent experiments. \**P* < 0.05, \*\**P* < 0.01, \*\*\*\**P* < 0.0001, two-tailed unpaired Student’s t test in (**d,h**), two-way ANOVA multiple-comparisons test in (**m**).

It has been shown that the strong tonic signaling of GD2 (14G2A) CAR stems from its unique FRs, which initiate CAR signaling independent of antigen-binding. Even when all complementary determining regions (CDRs) of GD2(14G2A) CAR were replaced by respective CDRs from CD19 CAR that showed little tonic signaling, the resulting CD19- CDR-GD2(14G2A)-FR chimeric CAR still shows constitutive tonic signaling ^51^. Inspired by this study, we replaced all CDRs of GD2 (14G2A) CAR by those of HER2 CAR (Fig. 7i,j), and the HER2-CDR-GD2 (14G2A)-FR chimeric CAR was tested for TIF cells induction (Fig. 7k). Compared with minimal induction of TIF program with native HER2 CAR (Fig. 7g,h), depletion of BCOR and ZC3H12A significantly enhanced the expansion of the chimeric HER2-CDR-GD2 (14G2A)-FR CAR-T cells, which persisted in vivo for at least 21 weeks (Fig. 7l,m). This CDRs swapping data demonstrates that: 1) the induction of GD2TIF cells does not require antigen recognition, since CDRs from either GD2(14G2A) CAR or HER2 CAR could work; and 2) combined with BCOR/ZC3H12A deficiency, tonic signaling is sufficient to induce TIF program when CAR target cells are not available, such as HER2 CAR-T cells in mice.

### Induction of human CAR-TIF cells

Finally, we tested whether TIF program could be induced in human T cells. We used a recently reported protocol wherein activated human T cells were transduced with lentivirus expressing both CAR and sgRNAs, followed by electroporation of Cas9 mRNA for gene editing (Fig. 8a)^55^. These CAR expressing (GFP^+^) and gene-edited human T cells were transferred into NSG mice for monitoring of CAR-T cell expansion and persistence (Fig. 8a and Extended Data Fig. 11). Similar to the induction of mouse GD2TIF cells (Extended Data Fig. 10), human GD2 CAR-T cells (CD4^+^ or CD8^+^) were only detected in sg*BCOR*/*ZC3H12A* group, but not other groups (Fig. 8b,c). These *BCOR*/*ZC3H12A* double-edited human GD2 CAR-T cells could expand and persist in secondary NSG mice upon serial transfer (Fig. 8d,e), demonstrating superior stemness of these cells, especially considering the exhaustion feature of this GD2 CAR ^51^. A significant portion of these serially transferred cells expressed both CD62L and PD-1 (Fig. 8e), a feature shared with mouse CAR19TIF, EGFRTIF and GD2TIF (Fig. 3 and Extended Data Figs. 9 and 10).

**Fig. 8.**
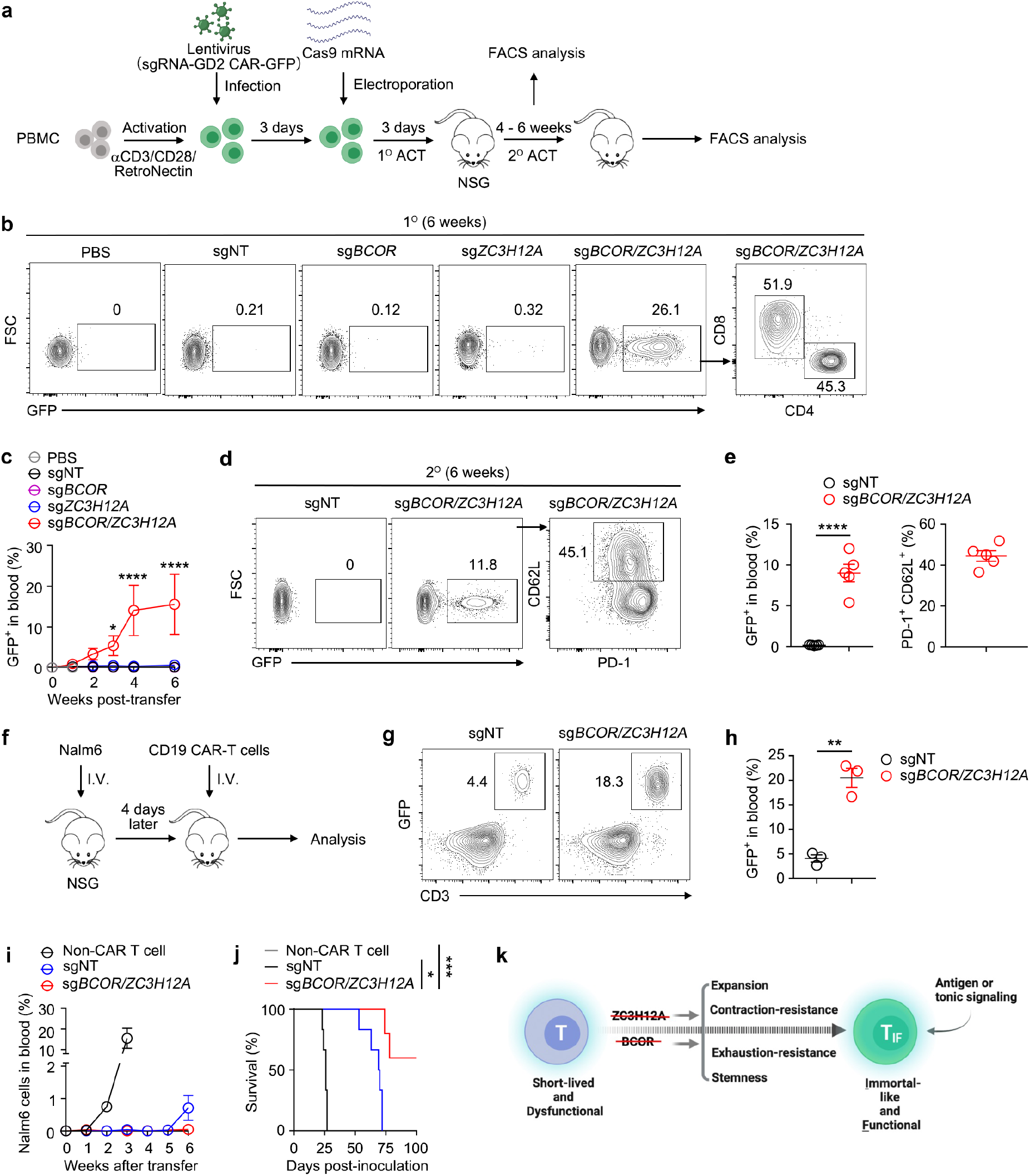
Induction CAR-TIF cells in human T cells. **a,** Experimental design for production of human CAR-T cells and editing of *BCOR* and *ZC3H12A*. **b,c**, Representative plots (**b**) and statistical analysis (**c**) of human GD2 CAR-T cells among single live cells in blood of 1° recipients at indicated time pointes are shown (n = 3 mice in PBS group, n = 5 mice in sgNT group, n = 5 mice in sg*ZC3H12A* group, n = 5 mice in sg*BCOR* group, n = 3 mice in sg*BCOR*/*ZC3H12A* group). **d,e**, Representative plots (**d**) and statistical analysis (**e**) of human GD2 CAR-T cells among single live cells in blood of 2° recipients 5 weeks post-transfer are shown (n = 5 mice for each group). **f**, Experimental design. Nalm6 cells (0.5 million) were intravenously injected into NSG mice. Four days later, 1.5 million of non-CAR T cells, sgNT or sg*BCOR*/*ZC3H12A* CD19 CAR-T cells were transferred into tumor-bearing mice intravenously. **g,h**, Representative plots (**g**) and statistical analysis (**h**) of CD19 CAR-T cells among single live cells in blood 6 weeks after transfer are shown (n = 3 mice for each group). **i,j**, Nalm6 in blood (**i**) and mice survival (**j**) were monitored (n = 6 mice in non-CAR T cell group, n = 6 mice in sgNT group, n = 5 mice in sg*BCOR*/*ZC3H12A* group). **k**, A model of CAR-TIF cells induction (see **Discussion**). Illustration created with Biorender.com. **c, e, h, i** and **j**, data represent mean ± SEM. Data are representative of three independent experiments. \**P* < 0.05, \*\**P* < 0.01, \*\*\**P* < 0.001, \*\*\*\**P* < 0.0001, two-way ANOVA multiple-comparisons test in (**c**), two-tailed unpaired Student’s t test in (**e,h**), log-rank (Mantel-Cox) test in (**j**).

We also tested the clinically approved CD19 CAR for human TIF cells induction ^56^. Since induction of CAR19TIF cells dependent on CD19^+^ target cells that is absent in NSG mice (Extended Data Fig. 4), we injected Nalm6 B cell leukemia cells into NSG mice as target cells, then transferred non-CAR T cells, control (sgNT) or *BCOR*/*ZC3H12A* double-edited human CD19 CAR-T cells (Fig. 8f). *BCOR*/*ZC3H12A* editing significantly enhanced the expansion/persistence of human CD19 CAR-T cells (Fig. 8g,h). Nalm6 cells grew quickly in NSG mice (Fig. 8i), which killed mice rapidly (Fig. 8j). The growth of Nalm6 cells in mice transferred with control CD19 CAR-T cells was repressed for 5 weeks, which re-emerged in blood starting from 6-week (Fig. 8i), followed by rapid death of these relapsed mice (Fig. 8j). However, mice received *BCOR*/*ZC3H12A* edited CD19 CAR-T cells repressed tumor cell growth and confer long-term survival of a large portion of recipient mice (Fig. 8i,j). These data demonstrate that TIF program could be induced in human T cells with clinically used CARs.

## Discussion

Generation of stem-like T cells for ACT has been considered the “holy grail” in T cell biology ^5^. We demonstrate that removal of two defined factors, ZC3H12A and BCOR, induces CAR-T cells into a novel state we called TIF, which exhibit unprecedented stemness but retain functionality of mature T cells.

Serial reconstitution of naive hosts is the golden standard to test stemness. For memory T cells, very few cells were recovered after 3 ∼ 4 generations of successive transfers ^11, 57, 58, 59^. Similarly, the repopulating capacity of HSCs decreases significantly after 3 to 4 transfers, which cease to repopulate between 4 to 6 transfers ^34, 35^. CAR19TIF cells were able to reconstitute naive hosts to a similar level of primary recipients after 6 successive transfers, which is unprecedented in known T cell subsets and even surpasses HSCs, suggesting CAR19TIF cells resemble iPSCs that self-renew infinitely ^36^. Importantly, repeated extensive expansion during serial transfers did not lead to exhaustion of CAR19TIF cells, which eradicated all target cells in recipients, demonstrating acquisition of ESCs/iPSCs-like self-renewing capacity by CAR19TIF cells is not at the cost of losing mature T cell function. To our best knowledge, the induction of CAR19TIF cells is the first example of reprogramming mature cells into such a state that can self-renew almost indefinitely like iPSCs but at the same time retain the functionality of mature T cells, which represents a novel existence of mammalian cells and reshapes our understanding of the relationship among stemness, differentiation and functionality ^20^. A recent study reported that, under appropriate vaccination protocols, virus-specific CD8^+^ T cells could be serially transferred in mice for 16 generations spanning 10 years ^60^, which were named as ISTCs. ISTCs were CD62L^neg^TIM-3^hi^, and thus phenotypically different from CAR-TIF cells that were CD62L^hi^TIM-3^neg^. Nevertheless, from data of this and our studies, it is emerging that T cells have the potential for maintaining infinite stemness without transformation.

CAR19TIF cells possess hybrid features of both stem and effector cells, as recently proposed for memory T cells ^61^. However, CAR19TIF cells differ from known T cell subsets in several aspects, especially their bona fide stemness. The simple and unique CD44^hi^CD62L^hi^PD-1^hi^ phenotype segregates CAR19TIF cells from known T cell subsets with stem-like properties, including TSCM (CD44^low^CD62L^hi^PD-1^neg^) ^6, 7^, TCM (CD44^hi^CD62L^hi^PD-1^neg^) ^62^ and TPEX (CD44^hi^CD62L^neg/low^PD-1^inter^) ^12, 13, 14, 15, 16^, since no known T cell subset co-expresses CD44, CD62L and PD-1 at high levels simultaneously. Previous studies showed that ZC3H12A-deficiency generated long-lived effector T cells and increased TCF1^+^ TPEX cells in CD19 CAR-T cells ^63, 64^. In our study, ZC3H12A- deficient CD19 CAR-T cells contracted rapidly after expansion, leaving no memory cells and B cell rebound ensued, demonstrating repressing ZC3H12A alone is unable to confer stemness. More importantly, ZC3H12A-deficiency was unable to expand T cell expressing EGFR or GD2 CAR in immunocompetent mice, demonstrating that the persistence-promoting effect of ZC3H12A-deficiency is context-dependent while the TIF program induced by BCOR- and ZC3H12A-double deficiency is conserved in CD19, EGFR and GD2 CAR-T cells.

Both ZC3H12A and BCOR are negative regulators of gene expression, which repress gene expression post-transcriptionally via promoting mRNA decay and epigenetically as transcription co-repressor ^33, 65^, respectively. Inhibition of these two proteins induced comprehensive changes of gene expression with thousands of genes up and down, which collectively reprogrammed CAR19TIF cells. Intriguingly, the expression of both ZC3H12A and BCOR is neither unique nor high in T cells ^65, 66^, so the successful induction of CAR19TIF cells by repressing these two proteins was unexpected. The exact mechanism of CAR19TIF cells reprogramming by the synergistic effect of ZC3H12A- and BCOR- double deficiency remains unclear. Although EOMES plays a critical role downstream of BCOR in CAR19TIF cells, other mechanisms must exist. Due to the inability of wild-type and BCOR-deficient CAR19T cells to expand in immunocompetent mice, we could not obtain these cells for comparation with CAR19TIF cells, which limits the power of mechanistic studies. Another unexplained phenomenon was that, unlike in CAR19T cells, ZC3H12A deficiency alone could not boost the expansion of GD2 and EGFR CAR-T cells in immunocompetent setting. Although BCOR deficiency also could not boost the expansion of GD2 and EGFR CAR-T cells, BCOR- and ZC3H12A-double deficiency induced GD2TIF and EGFRTIF exhibiting similar features with CAR19TIF cells. These data indicate that the synergy between BCOR and ZC3H12A is a must for the induction of CAR-TIF cells targeting solid tumor antigens. The underlying mechanisms are also difficult to dissect since we could not obtain BCOR- or ZC3H12A-deficient GD2 and EGFR CAR-T cells for side-by-side analysis. Future studies with appropriate models (e.g. infection) may solve this problem.

The induction and maintenance of CAR19TIF cells is antigen-dependent, while the induction of GD2TIF cells, likely EGFRTIF cells as well, depends on tonic signaling of CAR. Although CAR tonic signaling is usually considered detrimental to CAR-T cells ^51^, a recent study showed that it could promote CAR-T cells persistence under certain condition ^67^. TIF program is minimally induced in CARs with littler tonic signaling such as HER2 CAR (4D5) and GD2 CAR (K666). Importantly, HER2 CAR TIF cells could be induced when all their CDRs were placed in the unique FR of GD2 CAR (14G2A), which causes strong tonic signaling independent of antigen binding ^51^. Thus, the induction and maintenance of TIF program by ZC3H12A- and BCOR-double deficiency dependent on CAR signaling, either induced by antigen or tonic signaling (Fig. 8k). The facts that both GD2TIF and EGFRTIF cells could not survive in vitro, as well as their saturable niche in vivo, suggest that additional factor(s) contribute to the maintenance of these cells in vivo, which warrant further investigations.

Among hundreds of mice transferred with CAR-TIF cells in this study, none showed autoimmune or inflammatory response. Except for permanent absence of mature B cells in mice with CAR19TIF cells, mice with TIF cells were healthy with a normal endogenous T cell compartment. Deletion of ZC3H12A and BCOR in mature CD8^+^ T cells also did not lead to oncogenic transformation of TIF cells in any recipient mouse, which is consistent with their dormancy, inability to survive in vitro, saturable niche in NSG mice and downregulation of oncogene MYC. Further safety concerns about toxicity and oncogenic transformation can be solved by incorporation controllable CARs ^68^ or suicide genes ^69^ in CAR TIF cells.

Poor persistence of functional T cells is a common cause of cancer and chronic infection ^4, 5^. TIF cells exhibit bona fide stemness and unprecedented persistence in vivo, which will enhance current CAR-T cells therapy. Except for oncology, TIF cells may have other applications. First, CAR19TIF cells therapy do not require chemotherapeutic conditioning, thus they are more suitable than conventional CAR19T cells for autoimmune diseases currently treated by B cell depletion, which requires repeated dosing of depleting antibodies and has variable efficiency of B cell depletion among patients ^70^. Second, the large numbers of TIF cells persisting in vivo suggest that these cells, and their upgrades, may serve as a general platform to deliver therapeutic biologics requiring repeated dosing including antibodies, recombinant proteins, peptides and hormones.

## Methods

### Mice and cell lines

C57BL/6 (The Jackson Laboratory, Cat# JAX:000664, RRID:IMSR_JAX:000664), CD45.1 (The Jackson Laboratory, Cat# JAX:002014, RRID:IMSR_JAX:002014), *Rag1*-/- (The Jackson Laboratory, Cat# JAX:034159), NSG mice (The Jackson Laboratory, Cat# JAX: 005557, RRID: IMSR_JAX:005557) and Cas9 transgenic mice (The Jackson Laboratory, Cat# JAX:026430, RRID:IMSR_JAX:026430) were originally from The Jaxson Laboratory. Mice were maintained under specific pathogen-free conditions at the Laboratory Animal Research Center of Tsinghua University (Beijing, China). These animal facilities are approved by Beijing Administration Office of Laboratory Animal. All animal works were approved by Institutional Animal Care and Use Committee (IACUC). Age (6 - 12 weeks old)- and sex-matched mice were used for experiments.

To generate mCD19-expressing MC38 and B16F10 cell line, MC38 and B16F10 cells were transduced with retrovirus expressing mCD19 (pMIG-mCD19-IRES-GFP), GFP- and mCD19-double positive cells were sorted and expanded. To generate EGFR- expressing MC38 cell line, MC38 cells were transduced with lentivirus expressing EGFR (pLentiCas9-EGFR-T2A-Thy1.1), Thy1.1^+^ cells were sorted and expanded. Phoenix-Eco (ATCC, Cat# CRL-3214, RRID:CVCL_H717) and HEK293T cells (ATCC, Cat# CRL-3216, RRID:CVCL_0063) were cultured in DMEM (Gibco) containing 5% FBS, 2 mM glutamine, 100 units/ml penicillin and 100 µg/ml streptomycin in a humidified incubator at 37 °C. All cell lines were tested for Mycoplasma by the TransDectTM PCR Mycoplasma detection Kit (TRAN, Cat# FM311), and were confirmed to be negative.

### Vector and library construction

To generate a retroviral vector for expression of single guide RNA (sgRNA) together with a Thy1.1-P2A-CAR19 cassette, the hU6-sgRNA-EF1α-Cas9-P2A-puro expression cassette from lentiCRISPRv2 (Addgene, Cat# 52961) was cloned into pMSCV backbone (Addgene, Cat# 74056), then Cas9-P2A-puro was replaced by a Thy1.1 cassette. For anti-mCD19 CAR construct, the DNA sequence was obtained from a published report ^24^ and synthesized by Sangon Biotech (Shanghai). P2A-CAR19 constructs were cloned in frame with Thy1.1. This vector was named pMSCV-sgRNA-CAR19 (Fig. 1b), which was used for one-step generation of CAR19T cells with gene targeted by sgRNA after delivering into Cas9-expressing CD8^+^ T cells. The DNA sequence of anti-hCD19 CAR construct (47G-4) was from a published report ^56^ and synthesized by Sangon Biotech (Shanghai). The DNA sequence of anti-EGFR (Cetuximab) CAR, anti-GD2 CAR (14G2A and K666) and anti-HER2 (4D5) CAR were from published studies ^50, 51, 53, 54^ and synthesized by Sangon Biotech (Shanghai). The synthesized CAR cDNA was replaced in the pMSCV-U6-sgRNA-EFS-Thy1.1-P2A-CAR vector. The sequences of CAR-scFv are listed in Table S2.

For dual sgRNA screening, we inserted another U6 promoter to drive the second sgRNA (Fig. 1g). Then, the sgRNA part of Brie genome-wide sgRNA library (Addgene, Cat# 73632) for mouse was PCR-amplified and ligated into this vector via Gibson assembly (NEB, Cat# E2621S). The ligated product was precipitated, washed and electroporated into TOP10 bacteria which were sprayed on 40 plates. Plates were incubated at 33 °C for 15 hours, and bacteria clones were scratched off the plates for plasmid extraction. The coverage of this library was more than 200 colonies per sgRNA. The library was sequenced with Illumina NovaSeq 6000 (Berry Genomics). For screening, ∼ 150 million of CAR19T cells expressing dual sgRNAs were transferred into 50 B6 mice (3 million per mouse). On day 7 after transfer, CD8^+^Thy1.1^+^ CAR19T cells sorted from spleen of 10 recipient mice were pooled and used as input for normalization. Three months later, CD8^+^Thy1.1^+^ CAR19T cells were harvested from spleen of mice and pooled together for sgRNA enrichment analysis.

For triple gene knockout (Fig. 4h,i), we replaced the Thy1.1 marker of sg*Bcor*/*Zc3h12a* vector by GFP, and co-infected T cells with pMSCV-sgRNA-CAR19 with Thy1.1 marker. Thus, GFP^+^Thy1.1^+^ cells expressed three sgRNAs for triple knockout.

### Library construction for deep sequencing

To quantify the enrichment of sgRNA, genomic DNA was extracted by TIANamp Genomic DNA Kit (TIANGEN BIOTECH, Cat# DP304) according to manufacturer’s protocol. The sgRNA sequence was PCR-amplified using high-fidelity Q5 DNA polymerase (NEB, Cat# M0491L) with barcoded primers from genomic DNA for library construction, followed by deep sequencing (Illumina). The raw data of deep sequencing was trimmed to leave behind only sgRNA sequence with ENCoRE software. After comparison to the reference sgRNA sequence, the reads of each individual sgRNA in each sample were normalized within each sample as reads per million reads to offset the differences of sequencing depth among samples. For each gene, a *P* value was calculated by paired student t-test for examining the differences of gRNA abundance between input and end point for four gRNAs.

### Validation of gene editing in T cells expressing sgRNA

The editing of Bcor loci in CAR19TIF cells was examined by DNA sequencing. Briefly, CD8^+^Thy1.1^+^ and CD8^+^Thy1.1^-^ cells were sorted by S3e cell sorter (Bio-Rad) at 48 hours after spin-infection or one month after adoptive transfer. Genomic DNA were extracted for PCR amplification of genomic regions spanning the sgRNA cleavage sites. The amplified regions were sequenced to validate gene-editing. PCR primers were listed in Table S1.

### Western blot

The expression of ZC3H12A in control and ZC3H12A- and BCOR-double deficient CAR19TIF cells were examined by western blot. Four days after spin-infection, CD8^+^Thy1.1^+^ CAR19TIF cells and CD8^+^Thy1.1^-^ control cells were sorted by S3e cell sorter (Bio-Rad). Equal number of cells were lysed with Triton X-100 lysis buffer (40 mM Hepes, pH=7.4, 1% Triton X-100, 150 mM NaCl, 10 mM β-glycerol phosphate, 10 mM pyrophosphate, 2.5 mM MgCl2, 1X protease inhibitor) for 10 min on ice. The soluble fractions of cell lysates were isolated by centrifugation at 20,000 g at 4 °C for 10 min and quantified by a BCA kit (Thermo Fisher Scientific). Protein samples were denatured with the addition of 6 × SDS sampling buffer and incubated at 95 °C for 5 min. Protein samples were subjected to SDS-PAGE and immunoblotting analysis. Antibodies used to detect ZC3H12A and β-actin were listed in key resources table.

### Virus production

Retrovirus were packaged by co-transfection of Phoenix-Eco cells with indicated plasmid and helper plasmid pCL-Eco (Addgene, Cat# 12371) using calcium phosphate precipitate mediated transfection. The viral supernatant was collected at 48 and 72 hours post-transfection, filtered via 0.45 µM filters, aliquoted and frozen at -80 °C. Lentivirus were packaged by co-transfection of HEK293T cells with indicated plasmid encoding CAR, pMD2.G (Addgene, Cat# 12259) and psPAX2 (Addgene, Cat# 12260) using PEI transfection. Virial supernatant was collected at 48 and 72 hours post-transfection. The supernatants were filtered via 0.45 µM filters, and virus in supernatant were concentrated by ultracentrifugation (25,000 rpm for 2 hours). Concentrated virus were aliquoted and frozen at -80 °C.

### Primary T cells culture and infection

Mouse primary T cells were cultured in T cell medium (TCM): RPMI1640 medium (Gibco) supplemented with 5% fetal bovine serum (FBS), 2 mM glutamine, 55 µM β- mercaptoethanol, 1 mM sodium pyruvate, 100 units/ml penicillin, 100 µg/ml streptomycin and 2 ng/ml IL-2 in a humidified incubator at 37 °C with 5% CO2.

Single cell suspension were prepared from spleen and lymph nodes of Cas9-expressing mice (both male and female mice were used). CD8^+^ T cells were purified by a negative selection Kit (BioLegend, Cat# 480035), and purified CD8^+^ T cells were activated by 1 μg/ml anti-CD3 (BioXCell, Cat# BP0001-1, RRID:AB 1107634) and 1 μg/ml anti-CD28 (BioXCell, Cat# BP0015-1, RRID:AB 1107624) overnight.

Twenty-four hours after activation, viral transduction was performed by spin-infection with 2,000 g at 33 °C for 2 hours in the presence of 16 µg/ml polybrene (Sigma-Aldrich, Cat# H9268), followed by incubation for another 4 hours. Then, cells were washed and cultured in fresh TCM with IL-2. Twenty-four hours after spin-infection, the efficiency of transduction was determined by examination of reporter (Thy1.1 or GFP) positive cells by flow cytometry.

### Human CAR-T cell production and gene-editing

Peripheral blood mononuclear cells (PBMCs) were isolated from fresh whole blood of healthy donors by gradient centrifugation using Ficoll-Paque^TM^ PLUS (GE Healthcare, Cat# 17-1440-02). For T cell activation, 96-well plates were pre-coated with 5 μg/ml anti-hCD3 (BioLegend, Cat# 317326, RRID:AB_11150592), 1 μg/ml anti-hCD28 (Thermo Fisher Scientific, Cat# 16-0289-85, RRID:AB_468926) and 10 μg/ml RetroNectin (Takara, Cat# T100A). Then, PBMCs were loaded in these wells in human T cell media (X-VIVO media, Lonza, Cat# 04-418Q) supplemented with 5% human AB serum, 55 μM β- mercaptoethanol, 100 units/ml penicillin, 100 μg/ml streptomycin, 10 ng/ml hIL2, 10 ng/ml hIL7 and 10 ng/ml hIL15 in a humidified incubator at 37 °C with 5% CO2. After 24 hours, activated T cells were infected by lentivirus expressing CAR and sgRNA. Three days later, 2 million of CAR-T cells were electroporated with 1 μg Cas9 mRNA in 16-well Nucleocuvette™ Strips using EO-115 program (Lonza, per manufacturer’s instructions). Cells were expanded in human T cell media for another 3 days before analysis or transfer into mice.

The editing of *BCOR* and *ZC3H12A* genes in human CAR-T cells was examined by DNA sequencing. Briefly, human CAR-T cells were collected 3 days after electroporation. Genomic DNA were extracted for PCR amplification of genomic regions spanning the sgRNA cleavage sites. The amplified regions were sequenced to validate gene-editing. PCR primers were listed in Table S1.

### Adoptive T cell transfer

All adoptive transfers in this study were performed in the absence of any conditioning regimen. Twenty-four hours after spin-infection, indicated numbers of CD8^+^Thy1.1^+^ cells were transferred into age- and sex-matched B6 mice (Cas9^+^), *Rag1*^-/-^ mice, or NSG mice via tail vein. The presence of CD8^+^Thy1.1^+^ CAR-T cells and elimination of endogenous B cells (for CD19 CAR-T cells) in peripheral blood, spleen and indicated organs were examined by flow cytometry.

For serial transfer, ∼ 2 million of CAR19TIF cells from spleen of last batch of recipient mice were transferred into next batch of recipient mice via tail vein. The interval between transfers was one or two months as indicted in figures or figure legends. In some transfers, CAR19TIF cells were sorted by flow cytometry based on surface marker and CFSE (Thermo Fisher Scientific, Cat# C34554) signal, then re-labeled with CFSE for next transfer.

### Tumor models

In primary tumor protection model, MC38-mCD19 cells (2 × 10^5^) were subcutaneously injected into the right flank of male B6 mice. Two million CAR19TIF cells isolated from spleen of 2° donor mice (in 0.2 ml PBS) or PBS alone were injected into tumor bearing mice via tail vein. For memory protection model, MC38-mCD19 cells (5 × 10^5^) were subcutaneously injected into the right flank of male B6 mice previously transferred with 2 million of CAR19TIF cells (> one month) or PBS. Tumor size and mouse survival were recorded every 2 to 3 days. Tumor size was calculated by length × width. Mice bearing a tumor > 300 mm^2^ were considered as the endpoint of experiment and euthanized.

For lung metastasis model, B16F10-mCD19 cells (1 × 10^5^) were intravenously injected into B6 mice previously transferred with 2 million of CAR19TIF cells (> one month) or PBS. Three weeks after tumor injection, a cohort of mice were euthanized for macroscopic examination of tumor metastasis in the lungs. Another cohort of mice were monitored for survival.

For EGFR^+^ tumor model, MC38-EGFR cells (5 × 10^5^) were subcutaneously injected into the right flank *Rag1*^-/-^ mice. PBS, one million of sgNT or sg*Bcor*/*Zc3h12a* EGFR CAR-T cells were injected into tumor-bearing mice via tail vein. Tumor size and mouse survival were recorded every 2 to 3 days. Tumor size was calculated by length × width. Mice bearing a tumor > 300 mm^2^ were considered as the endpoint of experiment and euthanized.

For B cell leukemia model, Nalm6 cells (5 × 10^5^) were intravenously injected into NSG mice. Four days later, ∼1.5 million of untreated human T cells, sgNT CD19 CAR-T cells or sg*BCOR*/*ZC3H12A* CD19 CAR-T cells were injected into tumor-bearing mice via tail vein. The presence of CAR-T cells and Nalm6 cells in peripheral blood were examined by flow cytometry. Mouse survival was recorded every day.

### Flow cytometry

Single-cell suspension was prepared from blood, lymph nodes, spleen or indicated organs. Cell surface proteins were stained with indicated antibodies in the presence of Fc block in FACS buffer (PBS containing 1% FBS, 2 mM EDTA, 100 units/ml penicillin and 100 µg/ml streptomycin) at 4 °C for 15 min. Intracellular staining for cytoplasmic and nuclear proteins were performed with Transcription Factor Staining Buffer kit according to manufacturer’s instructions (BD Pharmingen). Dead cells were excluded by DAPI (BioLegend) staining or LIVE/DEAD Fixable Near-IR Dead Cell Stain Kit (Invitrogen). Antibodies for staining were from BD Pharmingen, BioLegend, or Invitrogen as listed with RRID. Samples were analyzed by LSRFortessa cytometer (BD). Flow cytometry data were analyzed using Flowjo software (https://www.flowjo.com). Cell sorting was performed on S3e cell sorter (Bio-Rad). All antibodies used in this study were listed in key resources table.

### In vitro killing assay

CAR19TIF cells were sorted from donor mice as effector cells. B cells were isolated from B6 mice as targeted cells. These two subsets were co-cultured in vitro at the indicated effector to target cell (E:T) ratios for 24 hours at 37 °C. In parallel, target cells were cultured alone to count basal cell number. The percentages of specific lysis for a given E:T ratio was calculated as (B cell number of control group - B cell number of CAR19TIF group)/B cell number of control group.

### In vitro stimulation assay

CAR19TIF cells and endogenous CD8^+^ T cells were stimulated with B cells (E/T:1/1) for 24 hours. Cell size and indicated protein expression were analyzed by flow cytometry.

### Metabolic assays

Oxygen consumption rate (OCR) and extracellular acidification rate (ECAR) were measured using Seahorse XFe96 analyzer following the manufacturer’s instructions. Briefly, endogenous CD8^+^ T and CAR19TIF cells were sorted and recovered in humidified incubator at 37 °C with 5% CO2 for 1 hour. Endogenous CD8^+^ T and CAR19TIF cells were suspended in XF medium and then plated in a poly-L-lysine-coated XF96 plate at the density of 3 x 10^5^ cells/well. OCR was measured in response to 2 μM oligomycin, 2 μM FCCP, 1 μM antimycin and 1 μM rotenone (Agilent, Cat# 103708-100) sequentially. ECAR was measured in response to 10 mM glucose, 4 μM oligomycin and 50 mM 2-DG (Agilent, Cat# 103020-100).

Untargeted metabolome analysis was performed using an UHPLC system (Vanquish, Thermo Fisher Scientific). In brief, endogenous CD8^+^ T cells and CAR19TIF cells were sorted by flow cytometry. Cells were stored in dry ice and sent to Biotree (Shanghai) for LC/MS analysis. The raw data were converted to the mzXML format using ProteoWizard and processed with an in-house program, which was developed using R and based on XCMS for peak detection, extraction, alignment and integration. Then, an in-house MS2 database (BiotreeDB) was applied in metabolite annotation. The cutoff for annotation was set at 0.3. KEGG (http://www.genome.jp/kegg/) and MetaboAnalyst (http://www.metaboanalyst.ca/) were used for pathway enrichment analysis.

Glucose uptake was examined by flow cytometry using 2-NBDG (MCE, Cat# HY-116215) in glucose-free medium (Macgene, Cat# CM15023). Cells were washed with glucose-free medium and incubated with 100 μM 2-NBDG in glucose-free medium at 37 °C for 30 min, and the uptake of 2-NBDG was measured by flow cytometry. To examine the effect of adenosine on 2-NBDG uptake, cells were preincubated with 100 μM ADORA2A inhibitor ZM241385 (MCE, Cat# HY-19532) for 24 hours. Then, the uptake of 2-NBDG was measured as described above.

### Bulk RNA sequencing and analysis

At indicated time points after transfer, Thy1.1^+^CD8^+^ CAR19TIF cells, EGFRTIF cells or endogenous CD8^+^ T cells were sorted by flow cytometry with purity > 95% from spleen of recipient mice. RNA samples were isolated and purified using TIANGEN RNAprep Pure Cell/Bacteria Kit, then shipped to BGI for library preparation and RNA sequencing on a DNBseq™. Raw FASTQ files from sequencing were aligned to reference genome and reference gene set using HISAT/Bowtie2. Differential gene analysis was performed by DEseq2(R). Genes were determined differentially expressed if FDR < 0.001 and log-fold change > 1 or < -1. GSEA and KEGG enrichment analysis with performed by ClusterProfiler (v3.14.0). Heat maps and volcano plots were plotted by using ggplots2.

### Single cell RNA sequencing (scRNA-seq) and analysis

Two months after transfer, CD8^+^Thy1.1^+^ CAR19TIF cells from recipient mice were sorted by flow cytometry and directly processed for scRNA-seq library preparation by using Chromium Single Cell 3′ Library& Gel Bead Kit v2 (10X Genomics) according to manufacturer’s protocol. The ultimate constructed and purified library with the target recovery of ∼ 6,000 single cells was sequenced on Illumina Hiseq. Cell Ranger toolkit (v2.0.0) was used for 10X Genomics scRNA-seq data alignment and quantification. The generated data files including aligned and filtered reads, barcodes and unique molecular identifiers were processed by Seurat (v3.1.1) for downstream analysis. For the data set, cells were considered as low-quality and then excluded if number of detected genes < 200 or > 4,000. Cells were also removed if their mitochondrial gene proportions were larger than 20%. Following normalization process, the top 2,000 variable genes were chosen for principle-component analysis. 1 to 10 PCs, determined by *JackStraw* function as significant ones, were selected for t-SNE and clustering analysis. Cluster specific genes were identified using *FindAllMarkers* (log FC threshold = 0.25) function. *FeaturePlot* and *VlnPlot* functions were also used for data visualization. For topic modeling analysis, *FitGoM* function from CountClust (v.1.12.0) was used with K number as 16 and tolerance value as 0.1. “Top genes” for a given topic were calculated by *ExtractTopFeatures* function.

### Quantifications and statistical analysis

The statistical information of each experiment, including the statistical methods, the *P* value and sample numbers (n) were shown in figure or figure legends. GraphPad Prism 8 (https://www.graphpad.com) was used to plot all graphs and to perform statistical and quantitative assessments. Error bars represent standard error of mean (SEM).

### Reporting summary

Further information on research design is available in the Nature Portfolio Reporting Summary linked to this article.

### Data availability

Sequencing datasets will be deposited in the Gene Expression Omnibus database later.

#### Acknowledgments

We thank Institute for Immunology at Tsinghua University for providing and maintaining equipment. We thank Xin Lin for providing plasmids containing EGFR sequence and EGFR CAR sequence; Xin Lin’s lab for technical help with human T cell culture and transduction; and Chen Dong’s lab for equipment. This research was supported by Vanke Special Fund for Public Health and Health Discipline Development (NO. 2022Z82WKJ013 to M.P.), National Natural Science Foundation of China (grant 31741085 to M.P. and 31800747 to N. Y.), Tsinghua University Initiative Scientific Research Program (2021Z to M.P.), Tsinghua University Initiative Scientific Research Program (to M.P.) and funds from Tsinghua-Peking Center for Life Sciences and Institute for Immunology at Tsinghua University (to M.P.).

## Author contributions

L.W., G.J. and Q.Z. performed experiments and analyzed the data; Y.L., Z.L., X.Z. and N.Y. provided technical help; N.Y. supervised the project; M.P. conceived and supervised the project, analyzed and interpreted data, and wrote the manuscript with inputs from all authors.

---

**Extended Data Fig. 1.**
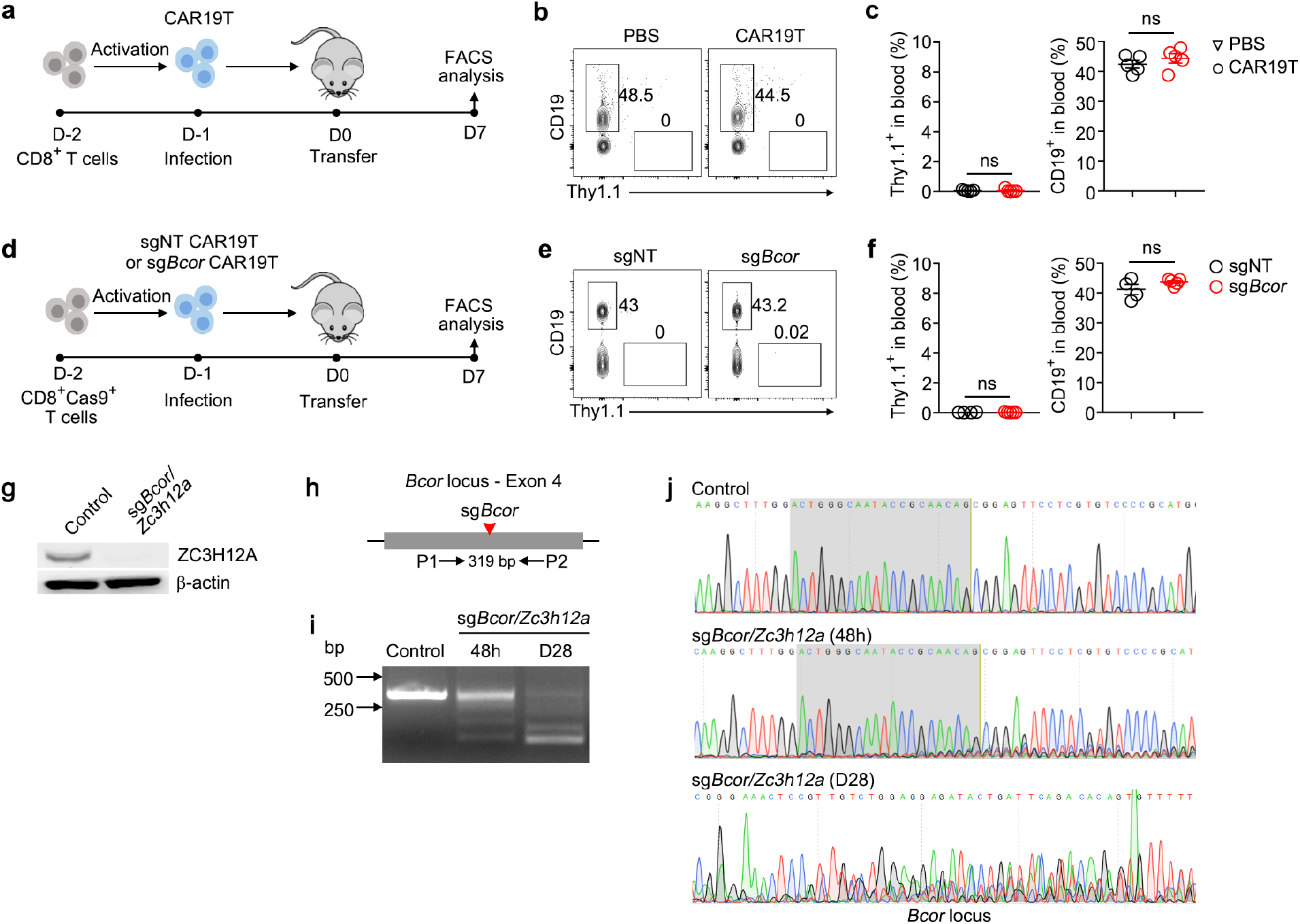
Wild-type and BCOR-deficient CAR19T cells do not expand in immunocompetent mice after adoptive transfer. **a,** Experimental design. CD8^+^ T cells were activated, transduced with retrovirus expressing CAR19 with a Thy1.1 marker. One million of CD8^+^Thy1.1^+^ CAR19T cells were transferred into B6 mice, PBS as control. After 7 days, Thy1.1^+^ CAR19T cells and CD19^+^ B cells from spleen were examined by flow cytometry. **b,c**, Representative plots (**b**) and statistical analysis (**c**) of Thy1.1^+^ CAR19T cells and CD19^+^ B cells among single live cells from spleen 7 days post-transfer are shown (n = 5 mice for each group). **d**, Experimental design for testing the effect of BCOR deficiency in CAR19T cells. **e,f**, Representative plots (**e**) and statistical analysis (**f**) of Thy1.1^+^ CAR19T cells expressing indicated sgRNA and CD19^+^ B cells among single live cells from spleen 7 days post-transfer are shown (n = 4 mice for each group). **g**, Immunoblot analysis of ZC3H12A expression in CD8^+^Thy1.1^-^ (control) and CD8^+^Thy1.1^+^ (sg*Bcor*/*Zc3h12a*) T cells 4 days post-transduction. β-actin was used as loading control. **h**, PCR detection of indels in *Bcor* loci. Red arrow indicates cleavage site of sg*Bcor*. A pair of PCR primers spanning cleavage site of Cas9/sg*Bcor* was used to amplify indicated genomic region. **i**, Editing of *Bcor* loci at 48 hours after transduction and 28 days post-transfer was examined by PCR. **j**, Editing of *Bcor* was examined by DNA sequencing. The PCR products from (**i**) were sequenced. Shade regions indicate sgRNA binding sites. Representative tracks of sequencing results are shown. **c,f**, data represent mean ± SEM from one of three independent experiments. ns, not significant, two-tailed unpaired Student’s t test. **g**-**j** data are representative of two independent experiments.

**Extended Data Fig. 2.**
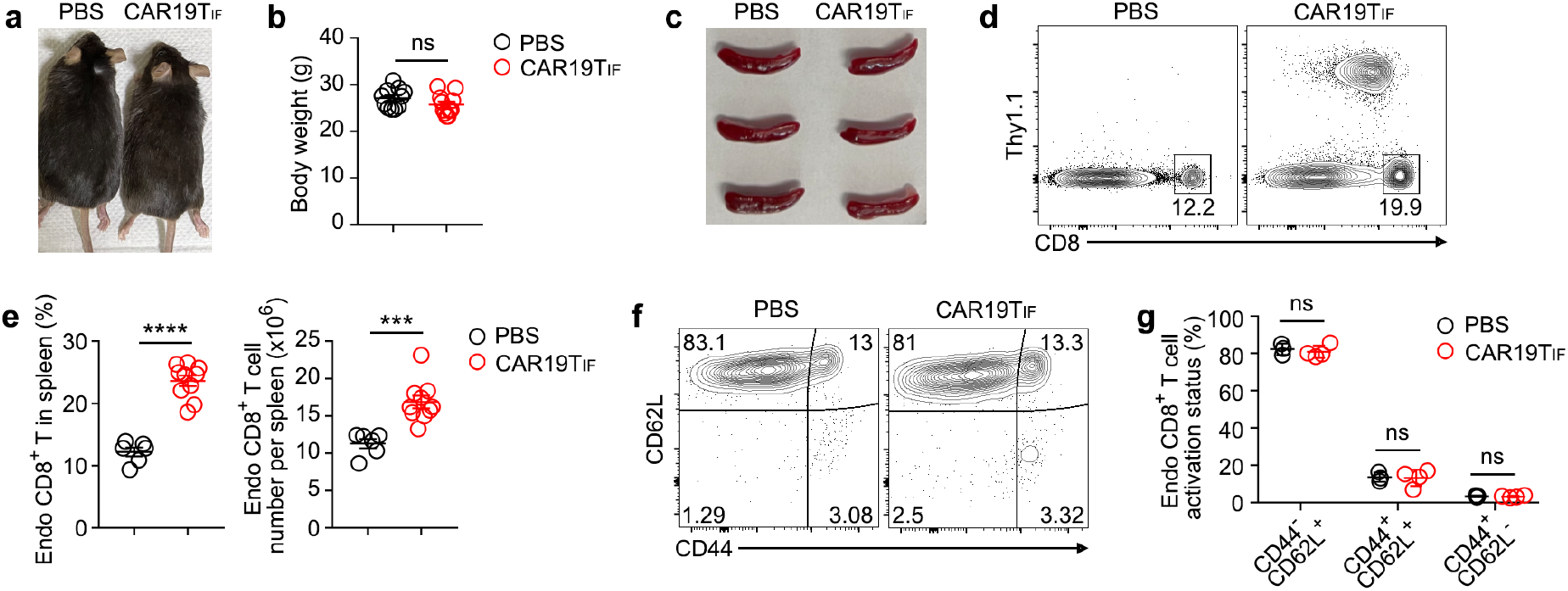
CAR19TIF cells persisting in vivo does not cause side-effects in mice. **a,** Representative images of mice six months after receiving CAR19TIF cells or PBS transfer. **b**, Body weight of mice six months after receiving CAR19TIF cells or PBS transfer (n = 11 mice for each group). **c**, Representative images of spleen from mice two months after receiving CAR19TIF cells or PBS transfer. **d,e**, Flow cytometry analysis of endogenous CD8^+^ T cells from spleen of mice two months after receiving CAR19TIF cells or PBS transfer. Representative plots (**d**) and statistical analysis (**e**) are shown (n = 6 mice in PBS group, n = 10 mice in CAR19TIF group). **f,g**, Flow cytometry analysis of activation status of endogenous CD8^+^ T cells from spleen of mice two months after receiving CAR19TIF cells or PBS transfer. Representative plots (**f**) and statistical analysis (**g**) are shown (n = 3 mice in PBS group, n = 4 mice in CAR19TIF group). **b, e** and **g**, data represent mean ± SEM. \*\*\**P* < 0.001, \*\*\*\**P* < 0.0001, ns, not significant, two-tailed unpaired Student’s t test.

**Extended Data Fig. 3.**
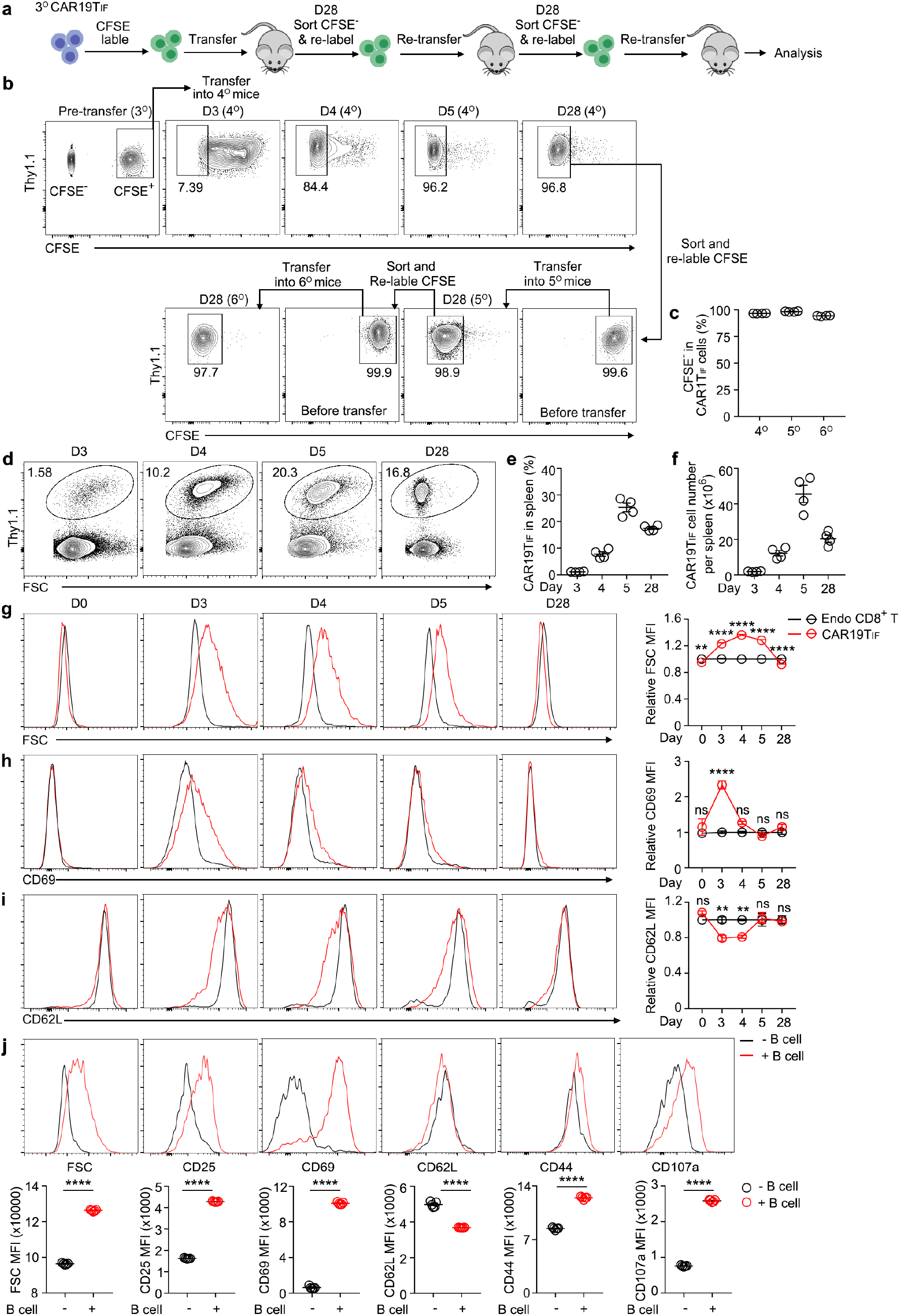
Phenotypical changes of CAR19TIF cells before and after expansion in recipient mice. **a,** Experimental design of serial transfer of carboxyfluorescein succinimidyl ester (CFSE)-labelled CAR19TIF cells. FACS sorted CAR19TIF cells were labelled with CFSE and transferred into B6 mice. **b,c**, Representative plots (**b**) and statistical analysis (**c**) of CFSE labelling and its dilution in CAR19TIF cells before and after transfer are shown (n = 4 mice for each group). **d-i**, Cell expansion (**d**-**f**) and phenotypical changes (**g**-**i**) were examined by flow cytometry at indicated days after transfer (n = 4 mice for each group). **j**, Flow cytometry analysis of phenotypic changes of CAR19TIF cells after co-culture with B cells. CAR19TIF cells isolated from spleen of 3° donor mice and B cells isolated from spleen of B6 mice were co-cultured at 1:1 ratio for 24 hours. The expression of indicated proteins was examined by flow cytometry (n = 5 replicates for each group). **c** and **e**-**j**, data represent mean ± SEM from one of three independent experiments. \*\**P* < 0.01, \*\*\*\**P* < 0.0001, ns, not significant, two-way ANOVA multiple-comparisons test in (**g**-**i**), two-tailed unpaired Student’s t test in (**j**).

**Extended Data Fig. 4.**
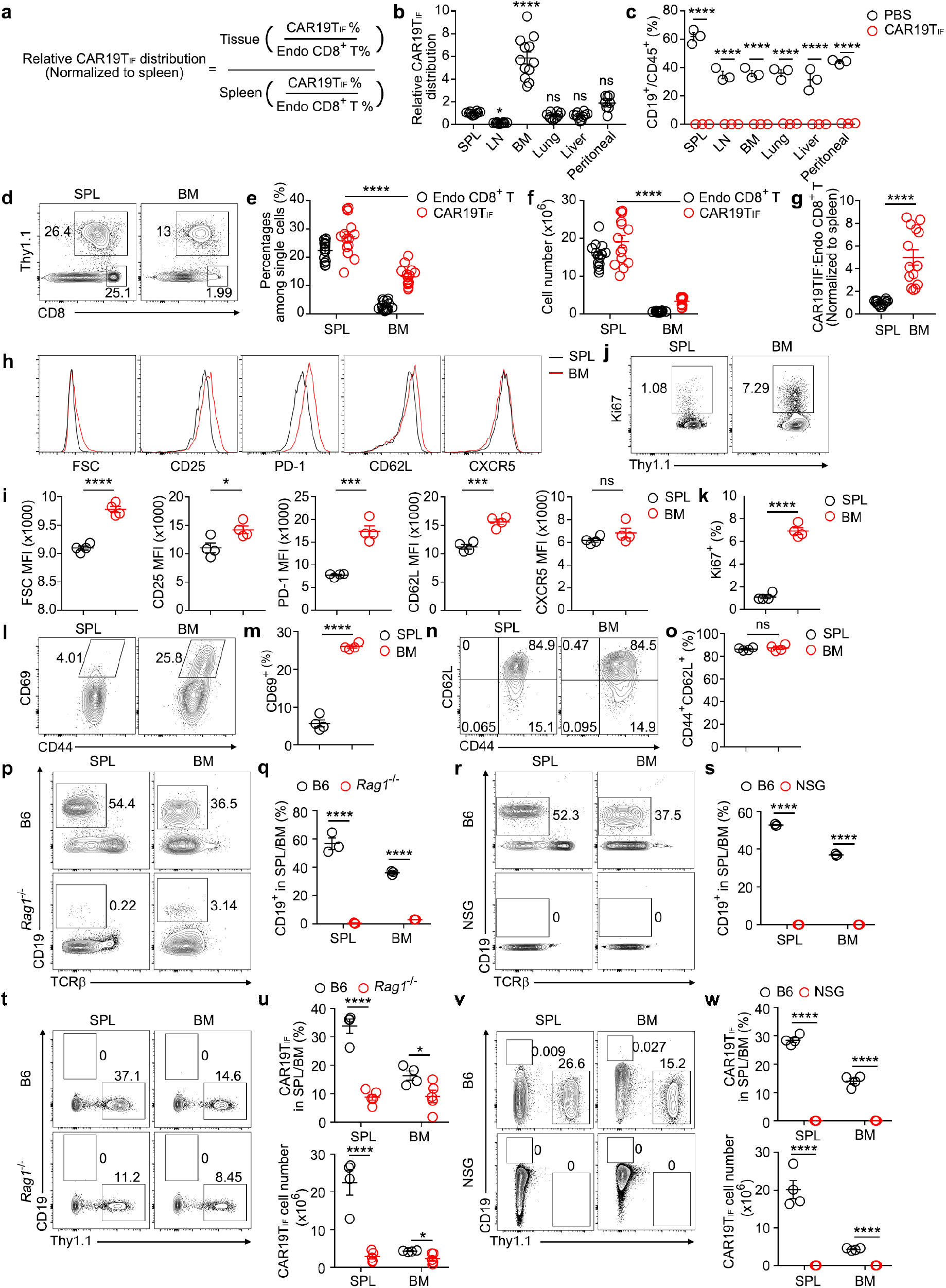
CAR19TIF cells are relatively enriched in BM and depedent on CD19^+^ cells. **a,** The formula used to calculate the relative distribution of CAR19TIF cells in different organs of B6 mice transferred with CAR19TIF cells. **b**, Relative distribution of CAR19TIF cells in spleen (SPL), lymph node (LN), bone marrow (BM), lung and peritoneal cavity (peritoneal) of recipient mice at one month post-transfer (n = 10 mice in SPL group, n = 10 mice in LN group, n = 12 mice in BM group, n = 10 mice in Lung group, n = 10 mice in Liver group, n = 8 mice in Peritoneal group for each group). **c**, Percentages of CD19^+^ B cells among CD45^+^ cells in indicted organs from mice at one month after transfer with CAR19TIF cells or PBS (n = 3 mice for each group). **d**-**f**, Percentages and cell numbers of Thy1.1^+^ CAR19TIF cells and endogenous CD8^+^ T cells from spleen and BM at one month post-transfer. Representative plots (**d**) and statistical analysis (**e,f**) are shown (n = 14 mice for each group). **g**, Normalized ratio of Thy1.1^+^ CAR19TIF cells to endogenous CD8^+^ T cells from spleen and BM (n = 14 mice for each group). **h,i**, Flow cytometry analysis of the expression of indicated protein on Thy1.1^+^ CAR19TIF cells from SPL and BM at one month post-transfer. Representative plots (**h**) and statistical analysis of mean florescence intensity (MFI) (**i**) are shown (n = 4 mice for each group). **j,k**, Percentages of Ki67^+^ CAR19TIF cells from spleen and BM at one month post-transfer. Representative plots (**j**) and statistical analysis (**k**) are shown (n = 4 mice for each group). **l,m**, Percentages of CD69^+^ CAR19TIF cells from spleen and BM of mice transferred with CAR19TIF cells at one month post-transfer. Representative plots (**l**) and statistical analysis (**m**) are shown (n = 4 mice for each group). **n,o**, Flow cytometry analysis of the percentages of CD44^hi^CD62L^hi^ cells among Thy1.1^+^ CAR19TIF cells from SPL and BM at one month post-transfer. Representative plots (**n**) and statistical analysis (**o**) are shown (n = 4 mice for each group). **p,q**, Flow cytometry analysis of CD19^+^ cells in spleen and BM of B6 and *Rag1*^-/-^ mice. Representative plots (**p**) and statistical analysis (**q**) are shown (n = 3 mice for each group). **r,s**, Flow cytometry analysis of CD19^+^ cells in spleen and BM of B6 and NSG mice. Representative plots (**r**) and statistical analysis (**s**) are shown (n = 3 mice for each group). **t,u**, Two million of Thy1.1^+^ CAR19TIF cells isolated from 2° donor mice were transferred into B6 or *Rag1*^-/-^ mice. After 4 weeks, percentages and cell numbers of CAR19TIF cells from spleen of recipient mice were examined by flow cytometry. Representative plots (**t**) and statistical analysis (**u**) are shown (n= 4 in B6 group, n = 5 mice in *Rag1*^-/-^ group). **v,w**, Two million of Thy1.1^+^ CAR19TIF cells isolated from 2° donor mice were transferred into B6 or NSG mice. After 4 weeks, percentages and cell numbers of CAR19TIF cells from spleen of recipient mice were examined by flow cytometry. Representative plots (**v**) and statistical analysis (**w**) are shown (n = 4 mice for each group). **b, c, e**-**g, i, k, m, o, q, s, u and w**, data represent mean ± SEM. \**P* < 0.05, \*\*\**P* < 0.001, \*\*\*\**P* < 0.0001, ns, not significant, one-way ANOVA multiple-comparisons test in (**b**), two-tailed unpaired Student’s t test in other panels.

**Extended Data Fig. 5.**
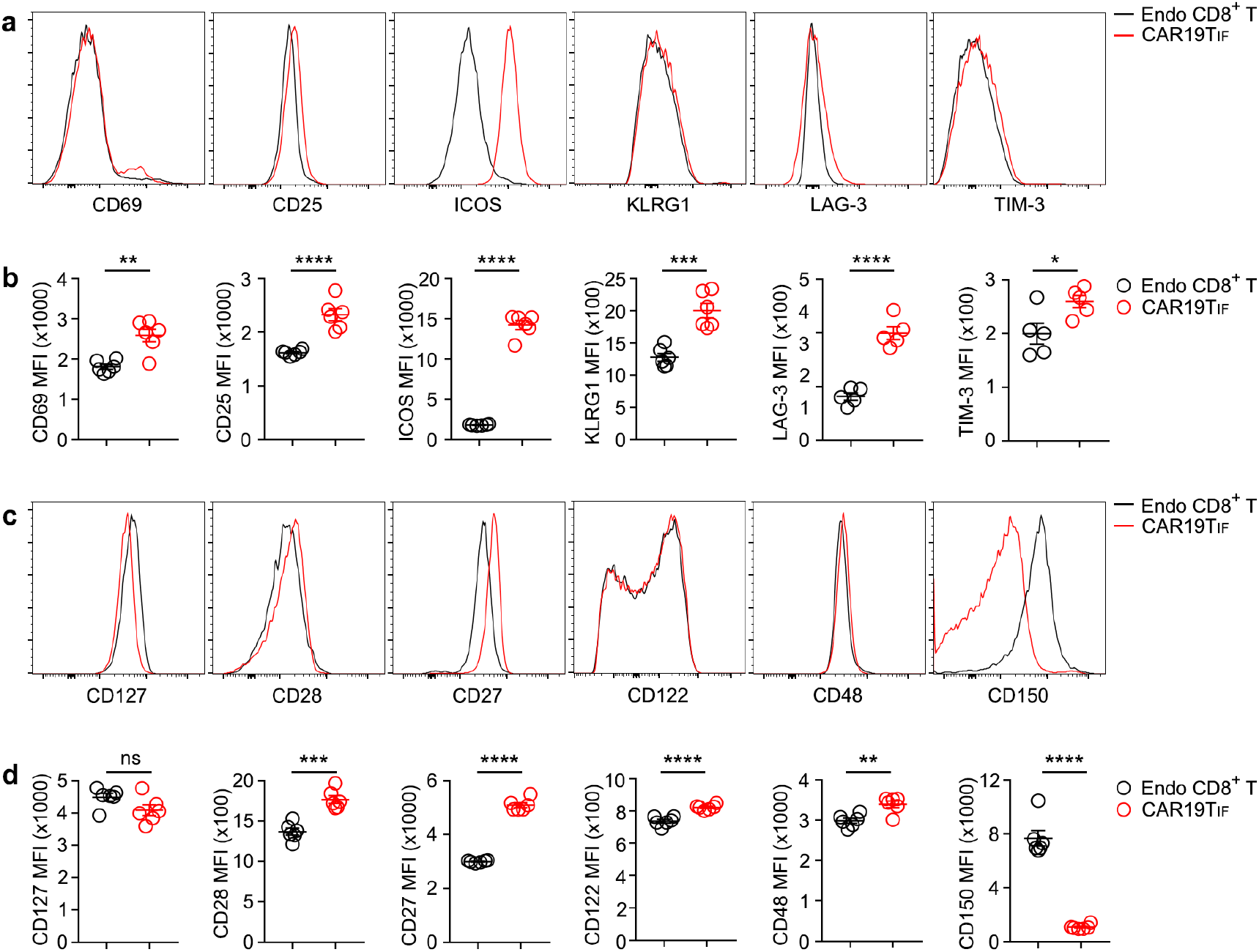
The expression of surface proteins by CAR19TIF cells. **a**-**d**, Flow cytometry analysis of the expression of indicated protein on endogenous CD8^+^ T cells and Thy1.1^+^ CAR19TIF cells isolated from spleen of 3° donor mice 2 months post-transfer. Representative plots (**a,c**) and statistical analysis of mean florescence intensity (MFI) (**b,d**) are shown (n = 5 - 6 mice for each group). **b** and **d**, data represent mean ± SEM. \**P* < 0.05, \*\**P* < 0.01, \*\*\**P* < 0.001, \*\*\*\**P* < 0.0001, ns, not significant, two-tailed unpaired Student’s t test.

**Extended Data Fig. 6.**
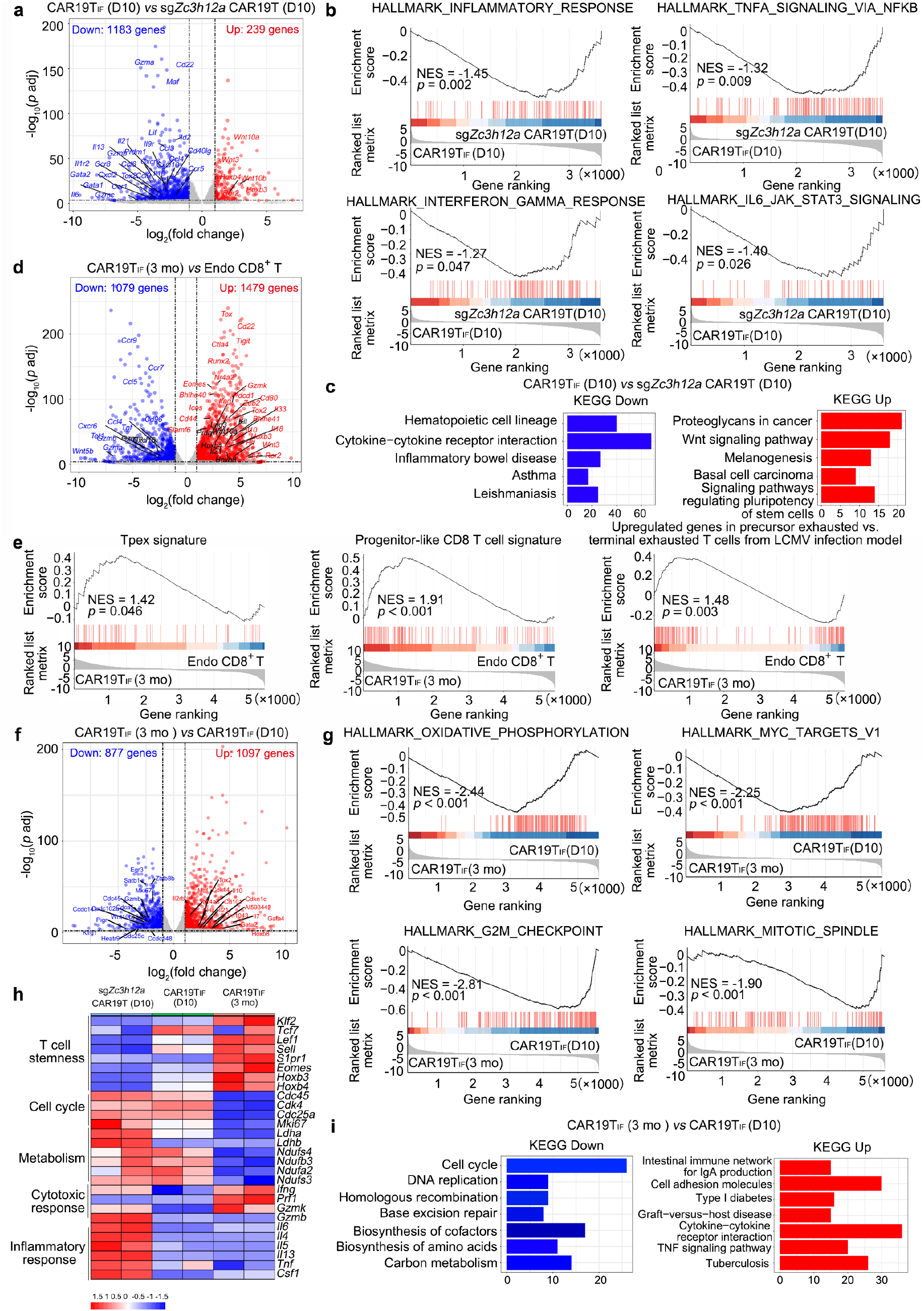
Bulk RNA-seq analysis reveals distinct but cooperative roles between ZC3H12A-deficiency and BCOR-deficiency during the reprogramming of CAR19TIF cells. **a,** A volcano plot of differentially expressed genes (DEGs) between ZC3H12A-deficient CAR19T cells and CAR19TIF cells 10 days post-transfer. **b**, GSEA plot of ZC3H12A-deficient CAR19T cells and CAR19TIF cells 10 days post-transfer. **c**, Pathway analysis of DEGs between ZC3H12A-deficient CAR19T cells and CAR19TIF cells 10 days post-transfer. **d**, A volcano plot of differentially expressed genes (DEGs) between CAR19TIF cells (3 months after transfer) and endogenous CD8^+^ T cells. **e**, GSEA plot of CAR19TIF cells (3 months after transfer) and Tpex from literatures. **f**, A volcano plot of differentially expressed genes (DEGs) of CAR19TIF cells between 10 days and 3 months post-transfer. **g**, GSEA of CAR19TIF cells at 10 days and 3 months post-transfer based on gene sets from MSigDB. **h**, A heatmap showing the expression of selected genes associated with T cell stemness, cell cycle, metabolism, cytotoxic function and cytokines in ZC3H12A-deficient CAR19T cells and CAR19TIF cells 10 days post-transfer, as well as CAR19TIF cells 3 months post-transfer. **i**, KEGG pathway analysis of up-regulated and down-regulated genes in CAR19TIF cells 3 months post-transfer vs 10 days post-transfer.

**Extended Data Fig. 7.**
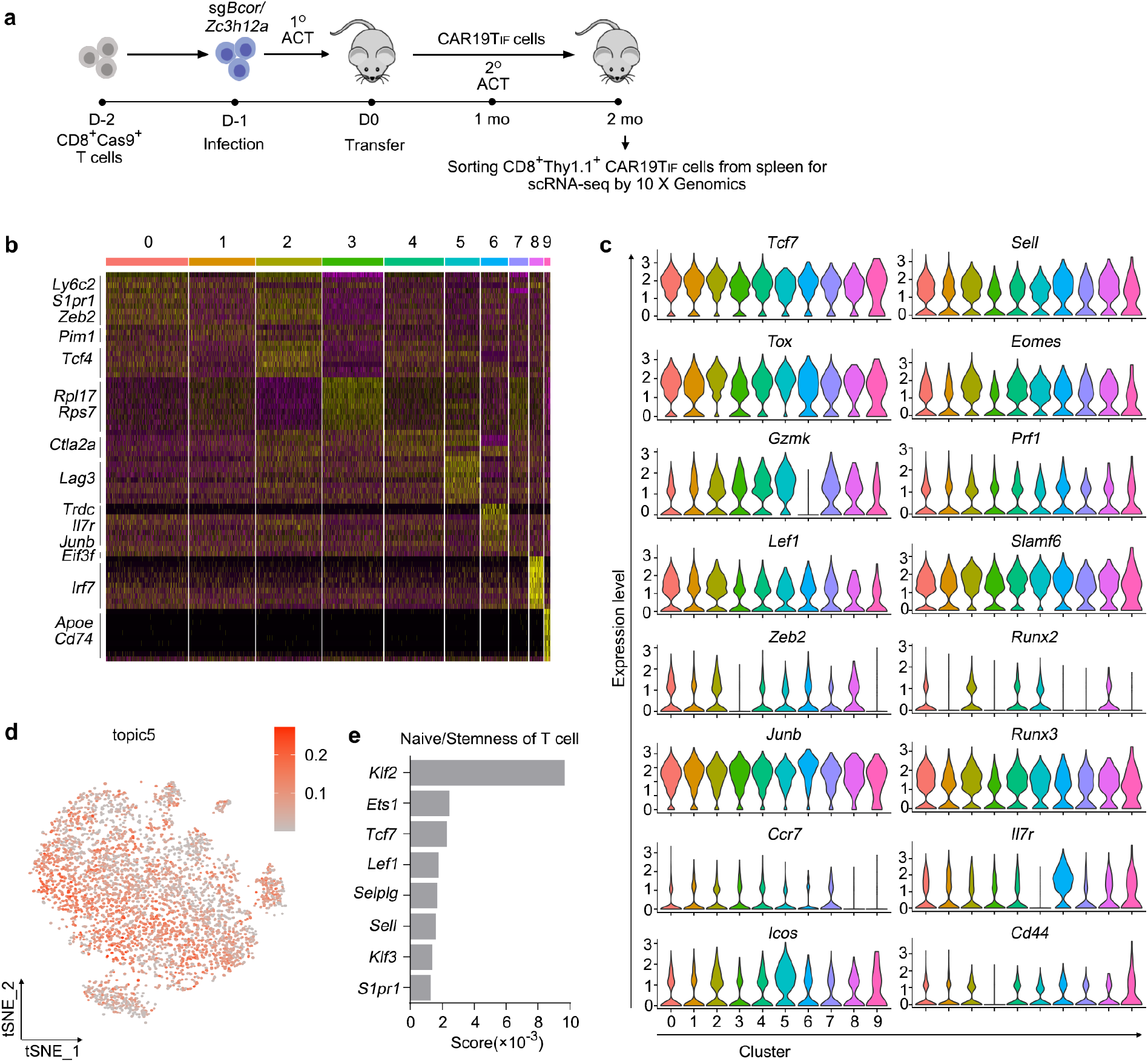
scRNA-seq analysis reveals homogeneity and hybrid features of CAR19TIF cells. **a,** Experimental setup for scRNA-seq. **b**, Heatmap visualization showing 10 clusters. Representative genes were labeled. **c**, Violin plots of relative expression of representative genes across all clusters. **d**, A two-dimensional tSNE plot of CAR19TIF cells scRNA-seq profiles colored by “topic5”’s weight in the cell. **e**, A bar plot of scores of “topic5” genes enriched and involved in T cell naïve/stemness state.

**Extended Data Fig. 8.**
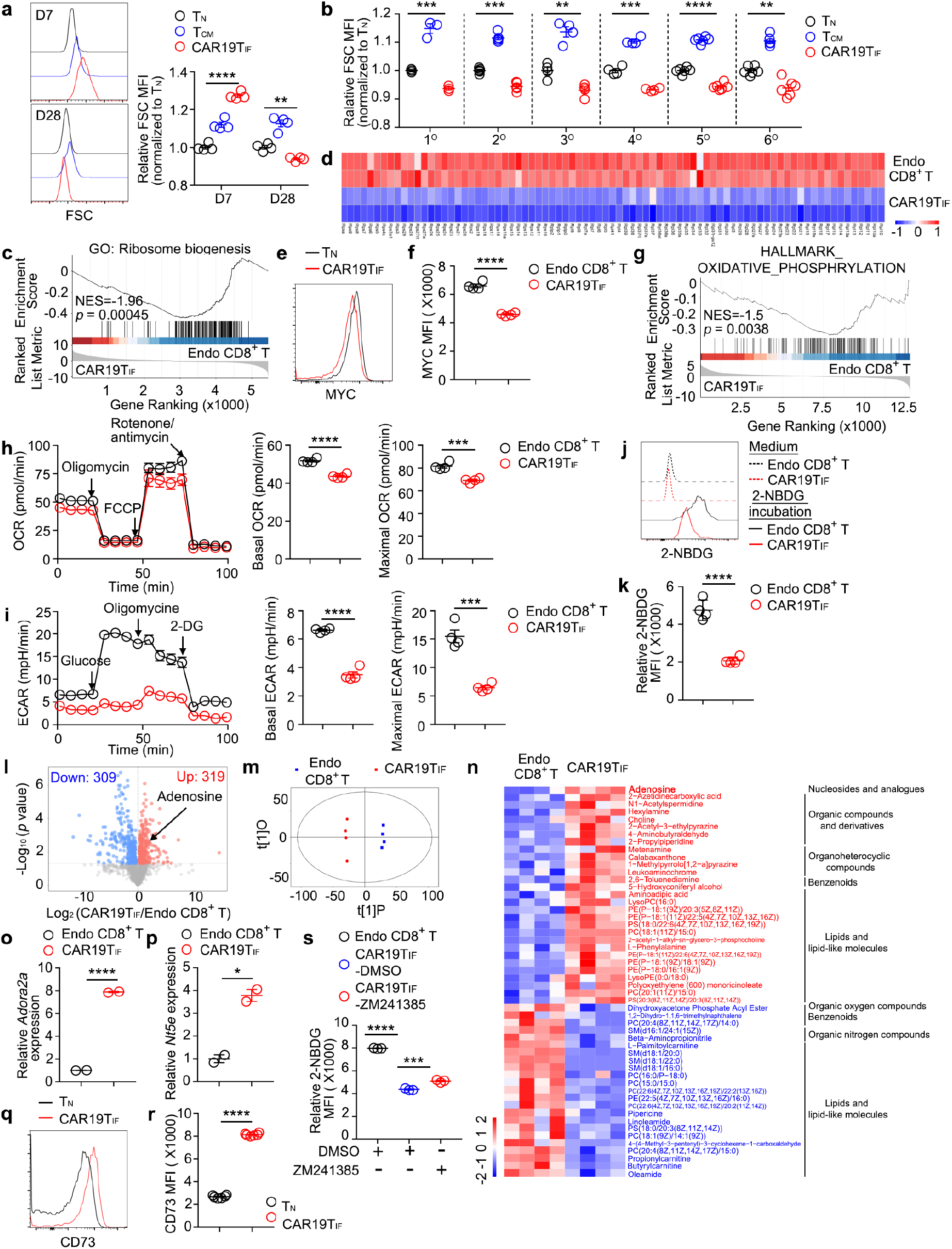
CAR19TIF cells are metabolically dormant in the absence of target cells. **a,** Flow cytometry analysis of splenic cell size 7 and 28 days post-transfer of CAR19TIF cells (n = 4 mice for each group). **b**, Flow cytometry analysis of cell size one month after transfer of CAR19TIF cells into different generations of recipients (n = 3 - 6 mice for each group). **c,** GSEA showing downregulation of genes involved in ribosome biogenesis in CAR19TIF cells (3 months post-transfer) compared with endogenous CD8^+^ T cells. **d**, A heatmap showing the expression of ribosome genes in endogenous CD8^+^ T cells and CAR19TIF cells (3 months post-transfer). **e,f**, Flow cytometry analysis of MYC expression in endogenous naive CD8^+^ T cells and CAR19TIF cells from spleen 28 days post-transfer. Representative plots (**e**) and statistical analysis of mean florescence intensity (MFI) (**f**) are shown (n = 4 mice for each group). **g**, GSEA showing downregulation of genes involved in oxidative phosphorylation in splenic CAR19TIF cells (3 months post-transfer) compared with endogenous CD8^+^ T cells. **h,i**, Seahorse experiments examining oxygen consumption rate (OCR) and extracellular acidification rate (ECAR) of endogenous CD8^+^ T cells and CAR19TIF cells sorted from spleen 28 days post-transfer. Representative plots and statistics are shown (n = 4 replicates for each group). **j,k**, Flow cytometry analysis of 2-NBDG uptake by endogenous CD8^+^ T cells and CAR19TIF cells from spleen 28 days post-transfer. Representative plots (**j**) and statistical analysis (**k**) are shown (n = 4 mice for each group). **l**, A volcano plot of differentially identified metabolites between endogenous CD8^+^ T cells and CAR19TIF cells from spleen of 2° donor mice.one month post-transfer by untargeted metabolomics. **m**, Metabolic feature of CAR19TIF cells and endogenous CD8^+^ T cells from spleen of 2° donor mice one month post-transfer. **n**, A heatmap showing the level of selected metabolites in endogenous CD8^+^ T cells and Thy1.1^+^ CAR19TIF cells from spleen. **o,p**, The expression of *Ador2a2* and *Nt5e* in endogenous CD8^+^ T cells and CAR19TIF cells (3 months post-transfer) from spleen. Data were from RNA-seq results (TPM). **q,r**, Flow cytometry analysis of CD73 expression on endogenous naïve CD8^+^ (TN) T cells and Thy1.1^+^ CAR19TIF cells from spleen of 5° donor mice one month post-transfer. Representative plots (**q**) and statistical analysis of mean florescence intensity (MFI) (**r**) are shown (n = 6 mice for each group). **s**, Endogenous CD8+ T cells and CAR19TIF cells from spleen of 5° donor mice one month post-transfer were pre-treated with ADORA2A inhibitor ZM241385 (100 μM) or DMSO for 24 hours, then incubated with 2-NBDG for another 30 min. 2- NBDG uptake was examined by flow cytometry (n = 3 mice for each group). **a,** **b, f, h, i, k, r** and **s**, data represent mean ± SEM from one of three independent experiments. \**P* < 0.05, \*\**P* < 0.01, \*\*\**P* < 0.001, \*\*\*\**P* < 0.0001, two-way ANOVA multiple-comparisons test in (**a**), one-way ANOVA multiple-comparisons test in (**s**), two-tailed unpaired Student’s t test in other panels.

**Extended Data Fig. 9.**
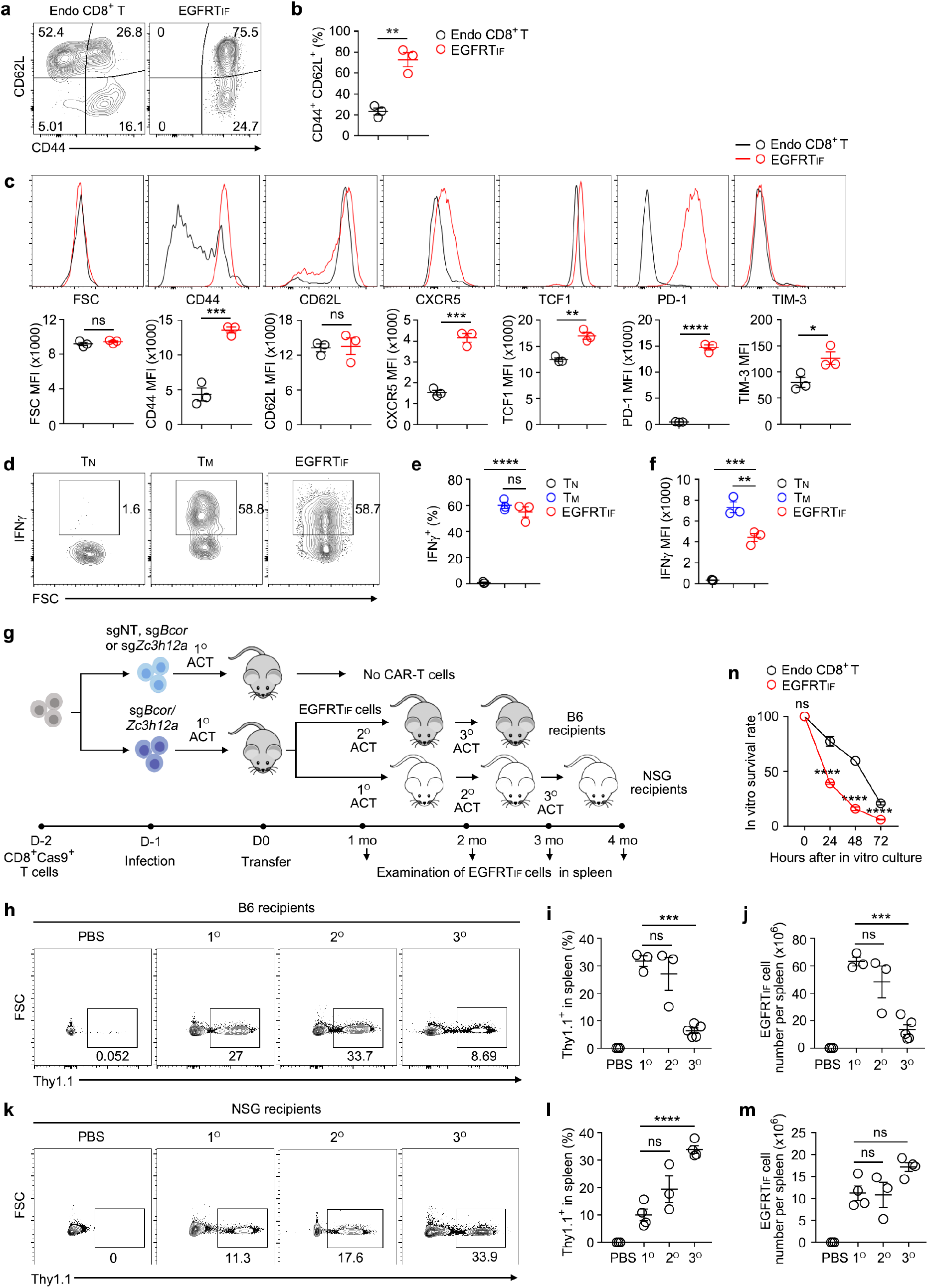
Characterization of EGFRTIF cells. **a**-**f**, Flow cytometry analysis of the expression of indicated protein on endogenous CD8^+^ T cells and EGFRTIF cells from spleen of primary recipient mice (4 weeks post-transfer). Representative plots (**a,c,d**) and statistical analysis (**b,c,e,f**) are shown (n = 3 mice for each group). **g**, Experimental design for serial transfer of EGFRTIF cells in B6 mice and NSG mice. **h**-**j**, Representative plots (**h**) and statistical analysis (**i,j**) of EGFRTIF cells among single live cells from spleen of B6 mice one month post-transfer are shown (n = 3 mice in PBS group, n = 3 mice in 1° group, n = 3 mice in 2° group, n = 5 mice in 3° group). **k**-**m**, Representative plots (**k**) and statistical analysis (**l,m**) of EGFRTIF cells among single live cells from spleen of NSG mice one month post-transfer are shown (n = 3 mice in PBS group, n = 4 mice in 1° group, n = 3 mice in 2° group, n = 4 mice in 3° group). **n**, Survival of EGFRTIF cells in vitro. EGFRTIF and endogenous CD8^+^ T cells were cultured in vitro in T cell medium for indicated hours (n = 6 replicates for each group). **b, c, e, f, i, j** and **l**-**n**, data represent mean ± SEM. Data are representative of three independent experiments. \**P* < 0.05, \*\**P* < 0.01, \*\*\**P* < 0.001, \*\*\*\**P* < 0.0001, ns, not significant, two-tailed unpaired Student’s t test in (**b,c**), two-way ANOVA multiple-comparisons in (**n**), one-way ANOVA multiple-comparisons test in other panels.

**Extended Data Fig. 10.**
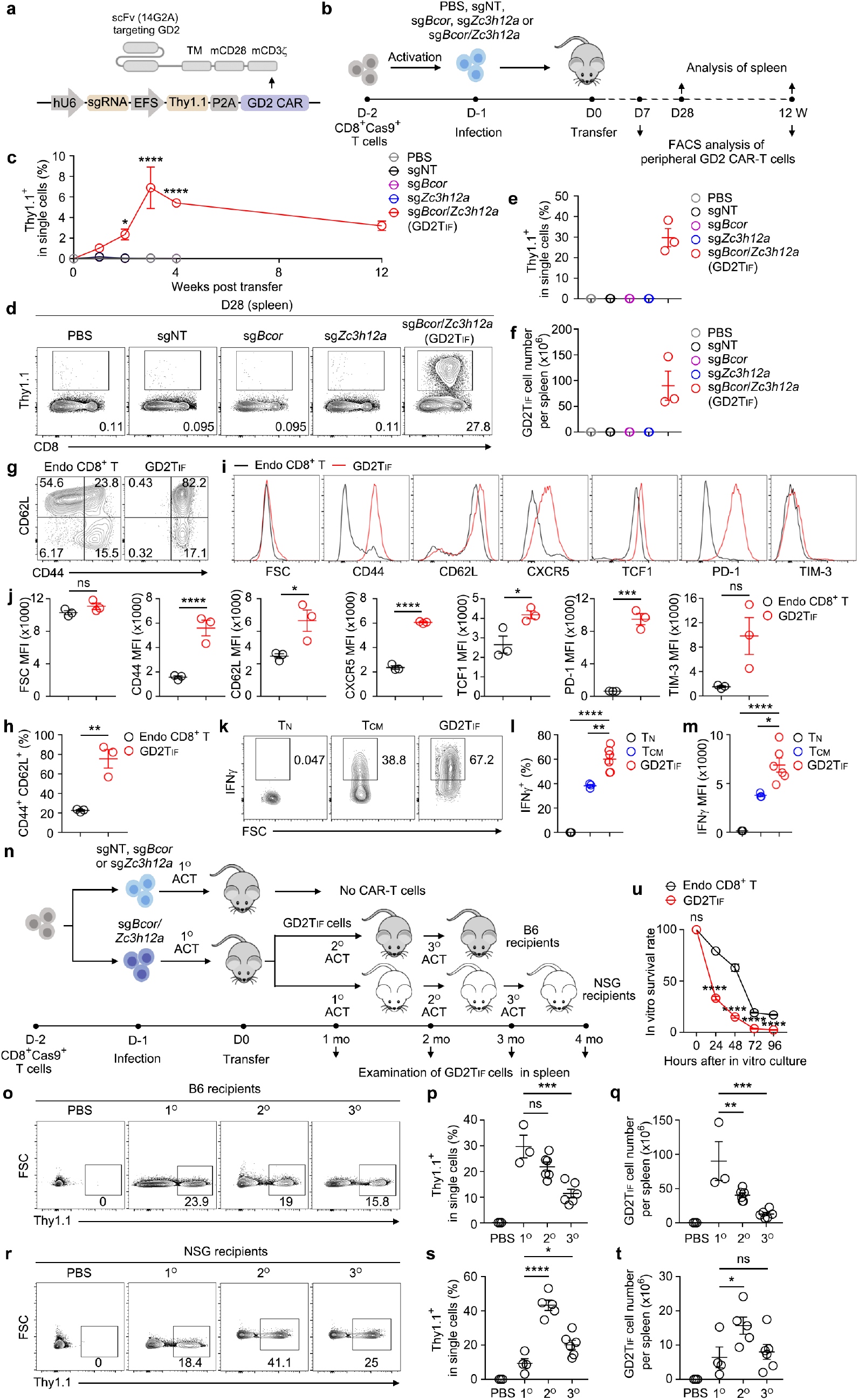
Induction and characterization of GD2TIF cells. **a,** The composition of anti-GD2 CAR with scFv derived from the 14G2A monoclonal antibody. **b**, Experimental design for the induction of GD2TIF cells. **c**, Statistical analysis of GD2TIF cells among single live cells from blood at indicated time (n = 3 mice for each group). **d**- **f**, Representative plots (**d**) and statistical analysis (**e,f**) of GD2TIF cells among single live cells in spleen 28 days post-transfer are shown (n = 3 mice for each group). **g**-**m**, Flow cytometry analysis of the expression of indicated protein on endogenous CD8^+^ T cells and GD2TIF cells from spleen of primary recipient mice 4 weeks post-transfer. Representative plots (**g,i,k**) and statistical analysis (**h,j,l,m**) are shown (n = 3 mice in **h,j**, n = 6 mice in **l,m**). **n**, Experimental design for serial transfer of GD2TIF cells in B6 mice and NSG mice. **o**-**q**, Representative plots (**o**) and statistical analysis (**p,q**) of GD2TIF cells among single live cells in spleen of B6 mice one month post-transfer are shown (n = 3 mice in PBS group, n = 3 mice in 1° group, n = 6 mice in 2° group, n = 6 mice in 3° group). **r**-**t**, Representative plots (**r**) and statistical analysis (**s,t**) of GD2TIF cells among single live cells in spleen of NSG mice one month post-transfer are shown (n = 3 mice in PBS group, n = 4 mice in 1° group, n = 5 mice in 2° group, n = 6 mice in 3° group). **u**, Survival of GD2TIF cells in vitro. GD2TIF and endogenous CD8^+^ T cells were cultured in vitro in T cell medium for indicated hours (n = 6 replicates for each group). **c-f, h, j, l, m, p, q and s-u**, data represent mean ± SEM. Data are representative of three independent experiments. \**P* < 0.05, \*\**P* < 0.01, \*\*\**P* < 0.001, \*\*\*\**P* < 0.0001, ns, not significant, two-way ANOVA multiple-comparisons test in (**c,u**), two-tailed unpaired Student’s t test in (**h,j**), one-way ANOVA multiple-comparisons test in other panels.

**Extended Data Fig. 11.**
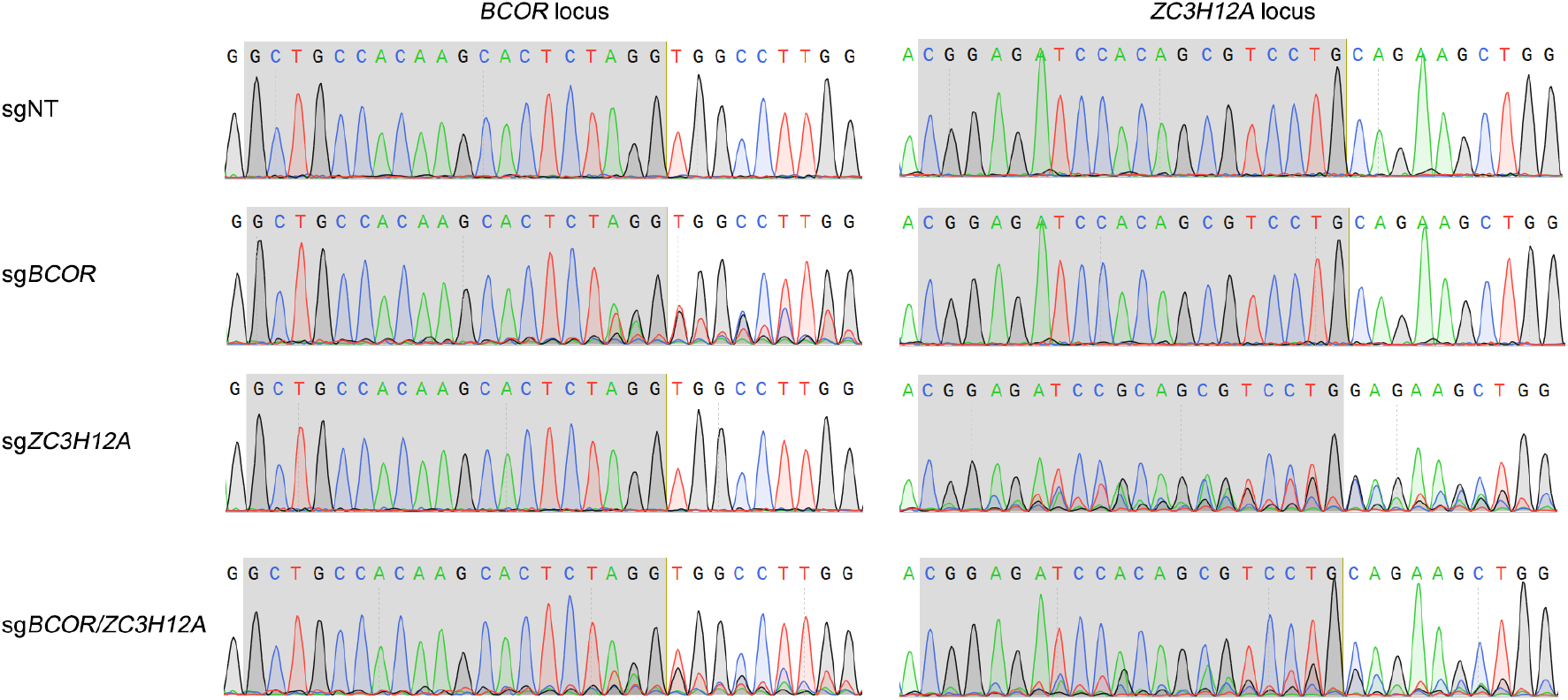
Examination of gene-editing in human T cells. Editing of *BCOR* and *ZC3H12A* loci in human T cells 72 hours after electroporation were examined by DNA sequencing. Shade regions indicate sgRNA binding sites. Representative tracks of sequencing results are shown.

**Supplementary Table 1.** Antibodies.

|  |  |  |
| --- | --- | --- |
| Anti-mouse CD19, biotin (MB19-1) | BioLegend | Cat# 101504,<br>RRID:AB_312823 |
| Anti-human CD19, APC (HIB19) | BioLegend | Cat# 302212,<br>RRID:AB_314242 |
| Anti-mouse Thy1.1, biotin (OX-7) | BioLegend | Cat# 202510,<br>RRID:AB_220141<br>7 |
| Anti-mouse Thy1.1, PE (OX-7) | BioLegend | Cat# 202524,<br>RRID:AB_159552<br>4 |
| Anti-mouse CD8α, APC (53-6.7) | BioLegend | Cat# 100712,<br>RRID:AB_312751 |
| Anti-mouse CD8α, eFluor 450 (53-6.7) | Thermo Fisher<br>Scientific | Cat# 48-0081-82,<br>RRID:AB_127219<br>8 |
| Anti-mouse CD8α, PE (53-6.7) | BioLegend | Cat# 100708,<br>RRID:AB_312747 |
| Anti-mouse CD8α, PE/Cyanine7 (53-6.7) | Thermo Fisher<br>Scientific | Cat# 25-0081-82,<br>RRID:AB_469584 |
| Anti-mouse CD8β, FITC (H35-17.2) | Thermo Fisher<br>Scientific | Cat# 11-0083-85,<br>RRID:AB_101729<br>46 |
| Anti-mouse CD3ε, biotin (145-2C11) | BioLegend | Cat# 100304,<br>RRID:AB_312669 |
| Anti-mouse CD44, PerCP/Cyanine5.5<br>(1M7) | BioLegend | Cat# 103032,<br>RRID:AB_207620<br>4 |
| Anti-mouse CD62L, PE/Cyanine7 (MEL-14) | BioLegend | Cat# 104418,<br>RRID:AB_313103 |
| Anti-mouse CD69, PE (H1.2F3) | Thermo Fisher<br>Scientific | Cat# 12-0691-83,<br>RRID:AB_465733 |
| Anti-mouse CXCR5, biotin (<br>L138D7) | BioLegend | Cat# 145510,<br>RRID:AB_256212<br>6 |
| Anti-mouse TCF1, PE (C63D9) | Cell Signaling<br>Technology | Cat# 14456,<br>RRID:AB_279848<br>3 |
| Anti-mouse c-Kit, FITC (2B8) | BD Pharmingen™ | Cat# 561680,<br>RRID:AB_109245<br>98 |
| Anti-mouse PD-1, FITC (29F.1A12) | BioLegend | Cat# 135214,<br>RRID:AB_106802<br>38 |
| Anti-mouse PD-1, PE (29F.1A12) | BioLegend | Cat# 135206,<br>RRID:AB_187723<br>1 |
| Anti-mouse TIM-3, PE/Cyanine7 (RMT3-23) | Thermo Fisher Scientific | Cat# 25-5870-82,<br>RRID:AB_257348<br>3 |
| Anti-mouse LAG-3, PerCP-eFluor 710 (C9B7W) | Thermo Fisher Scientific | Cat# 46-2231-82,<br>RRID:AB_111513<br>34 |
| Anti-mouse Eomes, Alexa Fluor 488 (Dan11mag) | Thermo Fisher Scientific | Cat# 53-4875-82,<br>RRID:AB_108542<br>66 |
| Anti-mouse Ki-67, FITC (SolA15) | Thermo Fisher Scientific | Cat# 11-5698-82,<br>RRID:AB_111513<br>30 |
| Anti-mouse TCRβ, Alexa Fluor® 647 (H57-597) | BioLegend | Cat# 109218,<br>RRID:AB_493346 |
| Anti-mouse CD25, FITC (PC61) | BioLegend | Cat# 102006,<br>RRID:AB_312855 |
| Anti-mouse CD107a, biotin (LAMP-1) | BioLegend | Cat# 121604,<br>RRID:AB_572003 |
| Anti-mouse ICOS, PE (C398.4A) | BioLegend | Cat# 313508,<br>RRID:AB_416332 |
| Anti-mouse KLRG1, APC (2F1) | BD Pharmingen™ | Cat# 561620,<br>RRID:AB_108957<br>98 |
| Anti-mouse CD127, FITC (SB/199) | BioLegend | Cat# 121106,<br>RRID:AB_493504 |
| Anti-mouse CD28, PE (37.51) | BioLegend | Cat# 102106,<br>RRID:AB_312871 |
| Anti-mouse CD27, APC (LG.3A10) | BD Pharmingen™ | Cat# 560691,<br>RRID:AB_172745<br>5 |
| Anti-mouse CD122, biotin (TM-β1) | BioLegend | Cat# 123206,<br>RRID:AB_940209 |
| Anti-mouse CD48, FITC (HM48-1) | BioLegend | Cat# 103404,<br>RRID:AB_313019 |
| Anti-mouse CD150, PE (TC1512F12.2) | BioLegend | Cat# 115904,<br>RRID:AB_313683 |
| Anti-mouse CD73, biotin (TY/11.8) | BioLegend | Cat# 127203,<br>RRID:AB_108906<br>3 |
| Anti-mouse CD45, APC-Cy™7 (30F11) | BD Pharmingen™ | Cat# 557659,<br>RRID:AB_396774 |
| Anti-mouse CD45.2, PE (104) | Thermo Fisher Scientific | Cat# 12-0454-83,<br>RRID:AB_465679 |
| Anti-mouse CD45.1, FITC (A20) | BioLegend | Cat# 110706,<br>RRID:AB_313495 |
| Anti-mouse IFNγ, PE (XMG1.2) | BioLegend | Cat# 505808,<br>RRID:AB_315402 |
| Anti-human CD8, biotin (RPA-T8) | BioLegend | Cat# 301004,<br>RRID:AB_314122 |
| Anti-human CD4, APC (OKT4) | BioLegend | Cat# 317416,<br>RRID:AB_571945 |
| Anti-human CD3, BV605 (SK7) | BD Pharmingen™ | Cat# 317416,<br>RRID:AB_271400<br>1 |
| Anti-human PD1, PE/Cyanine7 (A17188B) | BioLegend | Cat# 621615,<br>RRID:AB_283283<br>5 |
| Anti-human CD62L, PE (DREG-56) | Thermo Fisher Scientific | Cat# 12-0629-42,<br>RRID:AB_108041<br>44 |
| Anti-MCPIP1 (ZC3H12A) | Abcam | Cat# AB211659 |
| Anti-β actin | Santa Cruz | Cat# sc-47778,<br>RRID:AB_271418<br>9 |
| PE Streptavidin | BioLegend | Cat# 405204 |
| APC Streptavidin | BioLegend | Cat# 405243 |
| FITC Streptavidin | BioLegend | Cat# 405202 |
| APC/Cyanine7 Streptavidin | BioLegend | Cat# 405208 |
| Rabbit mAb anti-c-Myc/N-Myc, (D3N8F) | Cell Signaling Technology | Cat# 13987,<br>RRID:AB_263116<br>8 |
| Goat anti-Rabbit IgG (H+L), Alexa Fluor 647 | Thermo Fisher Scientific | Cat# A-21245 |
| DAPI | BioLegend | Cat# 422801 |
| LIVE/DEAD Fixable Near-IR Dead Cell Stain | Thermo Fisher Scientific | Cat# L34976 |

**Supplementary Table 2.**
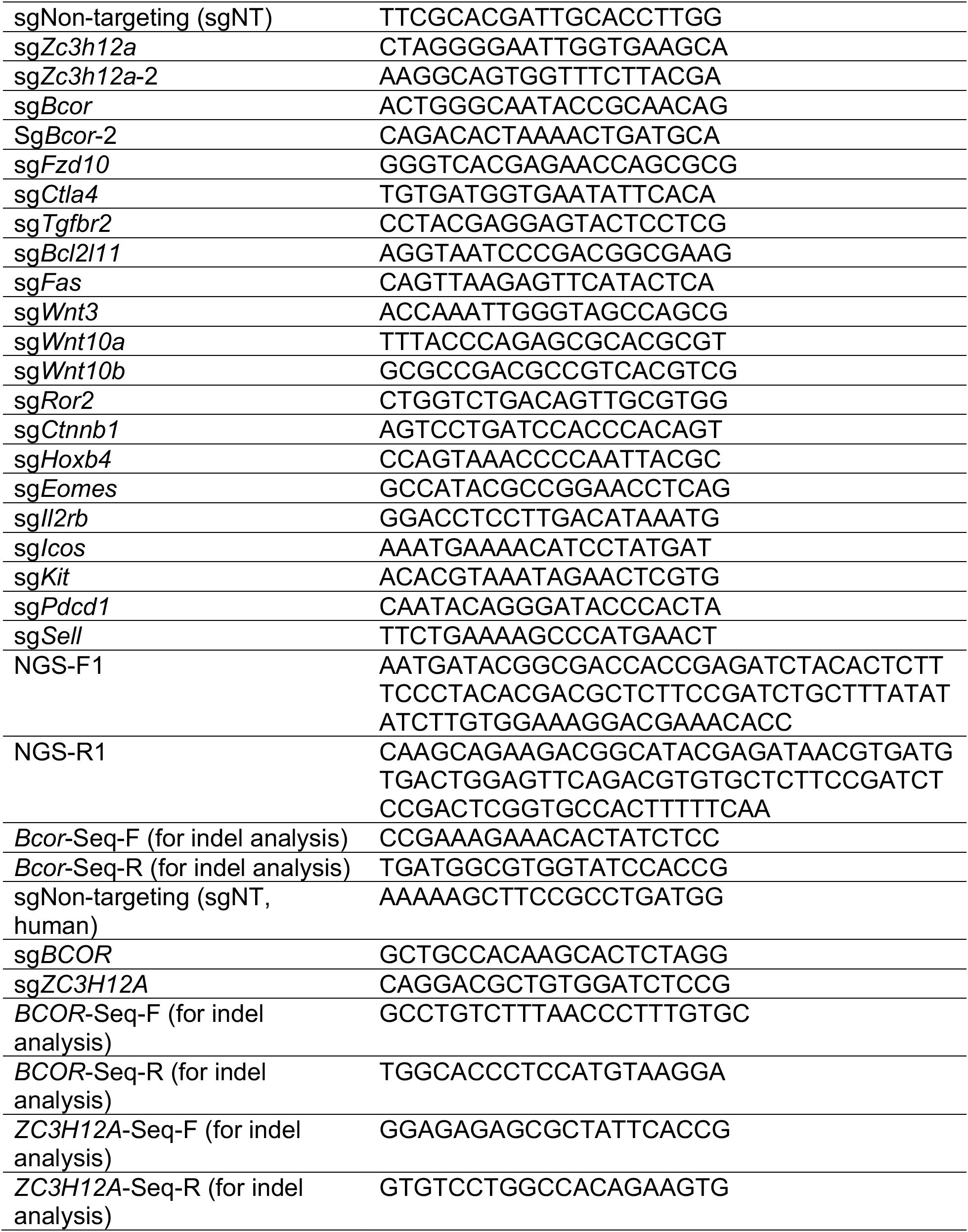
sgRNA sequences and primers.

**Supplementary Table 3.**
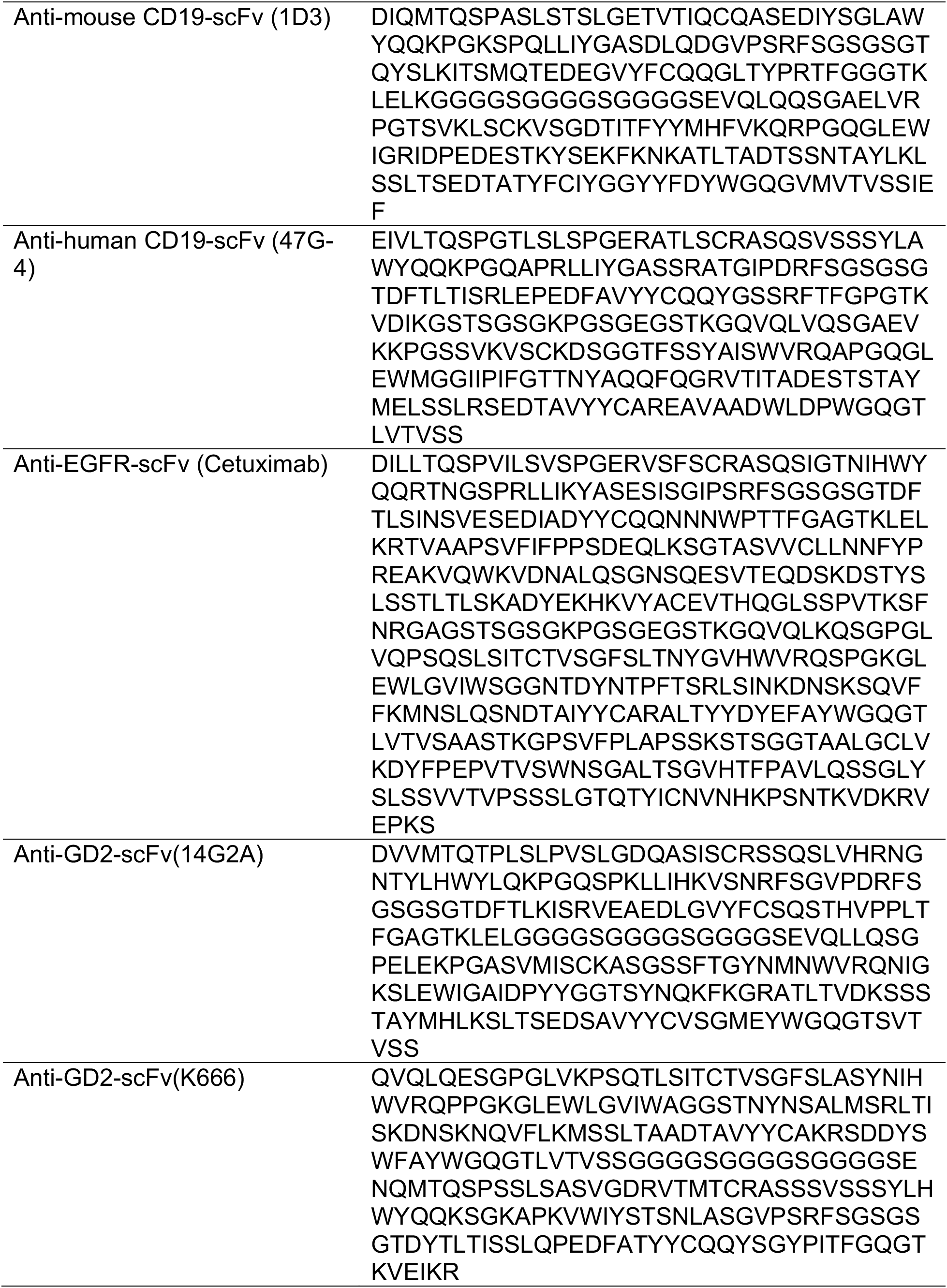
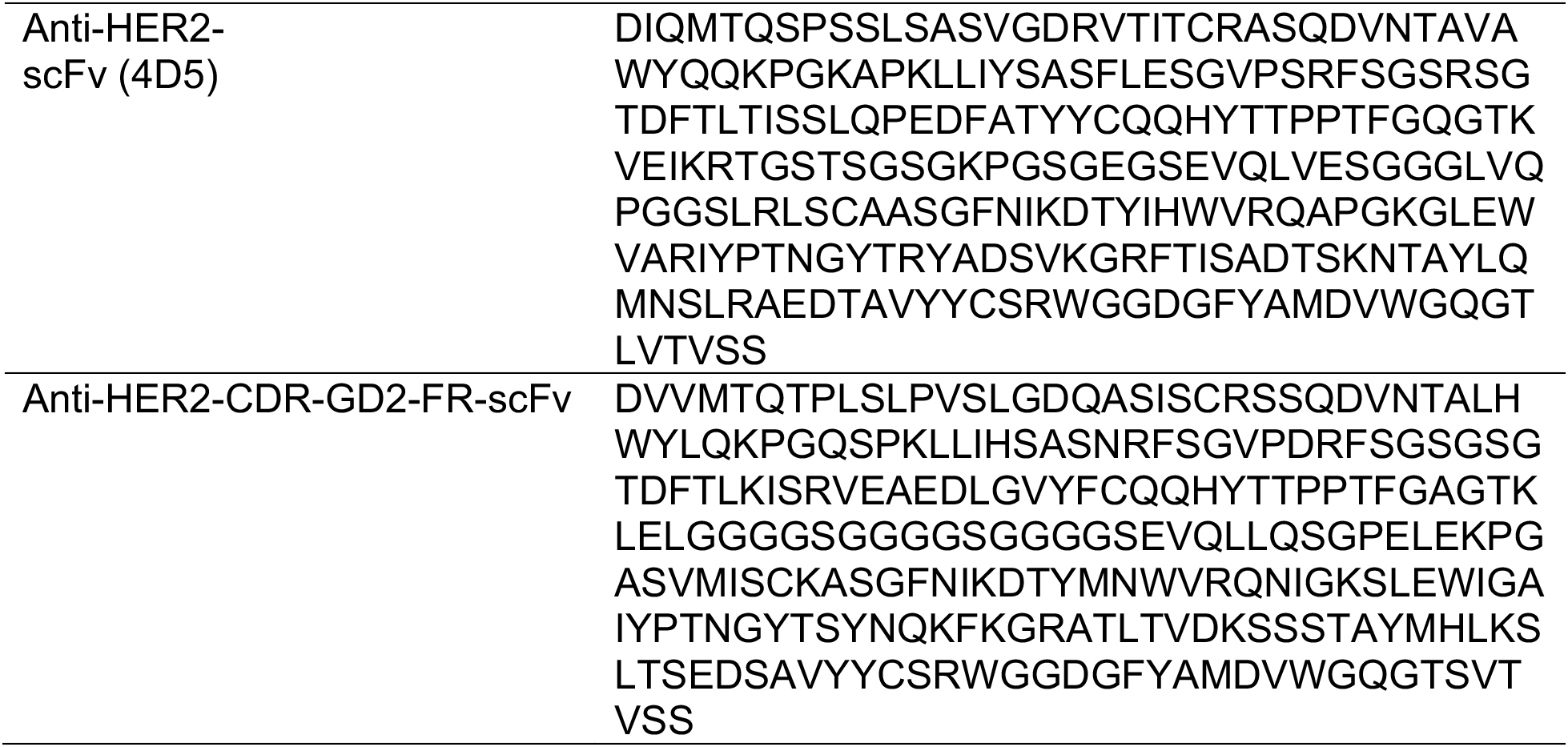
Sequences of CAR-scFv.

